# Reverse Genetics with a Full-length Infectious cDNA Clone of Bovine Torovirus

**DOI:** 10.1101/2020.10.28.358754

**Authors:** Ujike Makoto, Etoh Yuka, Urushiyama Naoya, Taguchi Fumihiro, Enjuanes Luis, Kamitani Wataru

## Abstract

Torovirus (ToV) has recently been classified in the new family Tobaniviridae, although it belonged to the Coronavirus (CoV) family historically. Reverse genetics systems for many CoVs have been established, but none exist for ToVs. Here, we describe a reverse genetics system using a full-length infectious cDNA clone of bovine ToV (BToV) in a bacterial artificial chromosome (BAC). Recombinant BToV containing genetic markers had the same phenotype as wild-type (wt) BToV. To generate two types of recombinant virus, the Hemagglutinin-esterase (HE) gene was manipulated, since cell-adapted wtBToV generally loses the full-length HE (HEf), resulting in soluble HE (HEs). First, recombinant viruses with HEf and HA-tagged HEf or HEs genes were rescued; these showed no significant differences in cell growth, suggesting that HE is not essential for viral growth in cells. Then, recombinant virus in which HE was replaced by the Enhanced Green Fluorescent Protein (EGFP) gene expressed EGFP in infected cells, but showed significantly reduced viral growth compared to wtBToV. Moreover, the recombinant virus readily deleted the EGFP gene after one passage. Interestingly, one variant with mutations in non-structural proteins (NSPs) showed improved EGFP expression and viral growth during serial passages, although it eventually deleted the EGFP gene, suggesting that these mutations contributed to EGFP gene acceptance. These recombinant viruses provide new insights regarding BToV and its reverse genetics will help advance understanding of this neglected pathogen.

**Importance:** ToVs are diarrhea-causing pathogens that have been detected in many species, including humans. BToV has spread worldwide, leading to economic losses. We developed the first reverse genetics system for Tobaniviridae using a BAC-based BToV. Using this system, we showed that recombinant BToVs with HEf and HEs showed no significant differences in cell growth. In contrast, clinical BToVs generally lose the HE gene after a few passages but some recombinant viruses retained the HE gene for up to 20 passages, suggesting some benefits of HE retention. The EGFP gene of the recombinant viruses was unstable and was rapidly deleted, likely via negative selection. Interestingly, one virus variant with mutations in NSPs was more stable, resulting in improved EGFP-expression and viral growth, suggesting that the mutations contributed to some acceptance of the exogenous EGFP gene without clear positive selection. The recombinant BToVs and reverse genetics developed here are powerful tools for understanding fundamental viral processes and their pathogenesis and for developing BToV vaccines.

## Introduction

Torovirus (ToV) belongs to order *Nidovirales*, family *Tobaniviridae*, subfamily *Torovirinae* and genus *Torovirus* (1). ToVs are a causative agent of diarrheic and respiratory diseases, and are detected in many species including bovine, equine, swine, goat, and human species (2–7). Although ToV infections are generally asymptomatic or do not have severe symptoms, the bovine torovirus (BToV) (8–16) and porcine ToV (PToV) (17–22) are distributed globally, causing economic loss. At present, there are no drugs and vaccines available for treatment and prevention of this disease.

ToVs are enveloped, positive-sense single-stranded RNA viruses with a genome 25-30kb in length, comprising the conserved six open reading frames. The first two-thirds of the genome contains ORF1a and ORF1b, with an overlap by a frameshift, encoding the replicase/transcriptase proteins (22–25). The remaining one-third of the genome encodes four structural proteins: spike (S), membrane (M), hemagglutinin-esterase (HE), and nucleocapsid (N) (26–30). ToV genome seemed to have occasionally undergone inter-genotype recombination events (31–33).

ToVs were difficult to propagate in cultured cells, with the exception of equine ToV (6); however, in the last 13 years, a new cell line, human rectal tumor-18 (HRT18) cells, has been shown to be susceptible to BToV, and several BToVs in Japan were successfully isolated and propagated using HRT18 cells (33–37). BToV with the full length HE gene was initially isolated from diarrheal feces, whereas cell-adapted BToV, after several passages in HRT18 cells, generally lost a full-length HE protein as a result of HE gene mutation (34, 35). This finding suggests that the HE protein is not essential for replication in cell culture and may instead suppress it.

ToVs historically belonged to the family *Coronaviridae*, which was divided into the subfamilies coronavirus (CoV) and ToVs, since both viruses are structurally and morphologically similar despite having some differences (24, 38–40). CoVs are mainly associated with respiratory and enteric diseases and are regarded as important pathogens in humans and animals. In human CoVs, in addition to four human CoVs causing mild upper respiratory disease (41), three novel life-threating CoVs that cause acute lung injury have emerged in the 21st century, namely, severe acute respiratory syndrome coronavirus (SARS-CoV) in 2002-2003 (42, 43), Middle East respiratory syndrome coronavirus (MERS-CoV) in 2012 (44), and SARS-CoV-2 in 2019 (45, 46). In animals, some CoVs cause severe or lethal diseases in swine (porcine epidemic diarrhea virus [PEDV] and transmissible gastroenteritis virus [TGEV]), bovine (bovine CoV), avian (infectious bronchitis virus [IBV]), mouse (mouse hepatitis virus [MHV]), and feline (Feline infectious peritonitis virus [FIPV]) animals. As such, numerous studies have been conducted on CoVs. Notably, reverse genetics systems for many human and animal CoVs, including the recently emerged SARS-CoV-2, have been established to study these fundamental viral processes and pathogenesis or to aid in vaccine development (47–68). Full-length cDNA-based reverse genetics systems of CoVs have been made, although there were obstacles such as large genome sizes, thereby complicating genome engineering, and the instability of some CoV replicase genes in bacteria. Currently, four systems are available: bacterial artificial chromosome (BAC) (48), in vitro ligation of cDNA fragments (47), vaccinia vector with full-length CoV cDNA (59), and the recently developed yeast artificial chromosome systems (60). Thanks to these systems, a greater understanding of CoVs has been gained.

In contrast to CoVs, only few studies of ToVs have been conducted, and no reverse genetics systems have been established since many ToVs are difficult to propagate in cells, and because less attention has been paid to ToVs as ToV infections are generally asymptomatic or not severe. In this study, we describe a reverse genetic system for BToV (Aichi strain) based on the cloning of a full-length genomic cDNA into BAC. Using this system, recombinant viruses with HA-tagged HE gene, untagged full-length HE gene, or with EGFP gene, were successfully rescued by manipulating the HE gene; these viruses were then characterized.

## Results

### Full-length genome sequence of wild-type (wt) BToV

To determine the parent BToV (Aichi strain) for reverse genetics, the full-length genomic sequences of three plaque-purified BToV were analyzed. We tried to determine the 5’ terminal end of these viruses using the 5’ RACE method, but several different sequences were observed in the terminal three bases; 5’-TG<u>ACGT</u>-3’, 5’-GT<u>ACGT</u>-3’, 5’-GG<u>ACGT</u>-3’, 5’-G<u>ACGT</u>-3’, 5’-<u>ACGT</u>-3’, 5’-<u>(-)CGT</u>-3’. The sequence at the 5’ terminal end of three published BToVs (25, 33) and two published PToVs (Accession # JQ860350 and KM403390) all shared the 5’-TG<u>ACGT</u>-3’ sequence, whereas equine ToV showed 5’-<u>ACGT</u>-3’ (74). Whether the differences observed in the BToVs were due to the intrinsic features of BToV or due to suboptimal experimental condition of the 5’ RACE method remained unclear. However, 5’-TG<u>ACGT</u>-3, previously reported in published BToVs and PToVs, was found in only one out of seven in our analysis, whereas 5’-<u>ACGT</u>-3’ reported in equine BToV is a consensus sequence. Thus, in this study, the sequence at 5’ terminal end of BToV was set as 5’-<u>ACGT</u>-3’. Two BToV clones showed the same sequence; from these, one was selected as the parent virus (wtBToV).

### Construction of a full-length BToV genome in BAC

Full-length BToV (Aichi strain) genome cDNA was assembled into the pBeloBAC11 plasmid carrying cytomegalovirus (CMV) immediate-early promoter, hepatitis delta virus ribozyme (Rz), and bovine growth hormone (BGH) termination, and the cDNA was inserted downstream off the CMV promoter and flanked by a 25-poly(A) sequence. The BToV genome cDNA was amplified into eight PCR fragments (BToV-A to H) carrying homology arms (hms) at both terminals, and these were sequentially assembled (Fig. 1A) using the Red/ET recombination method described in the Materials and Methods (Fig. 1B). After assembly of the eight fragments, the BAC carrying the full-length BToV genome cDNA (pBAC-BToV^mut1^) contained two single mutations at nucleotide (nt) G3399Tand at nt T8469G, and the 1,350 bp *E. coli* chromosomal derived sequence was inserted at nt 10,136, likely to counter the toxic regions in bacteria observed in some CoVs (47, 62, 63) (Fig. 2A). These two single mutations and *E. coli* sequence were reverted as described in the Materials and Methods (Fig. 2A).

**Fig. 1.**
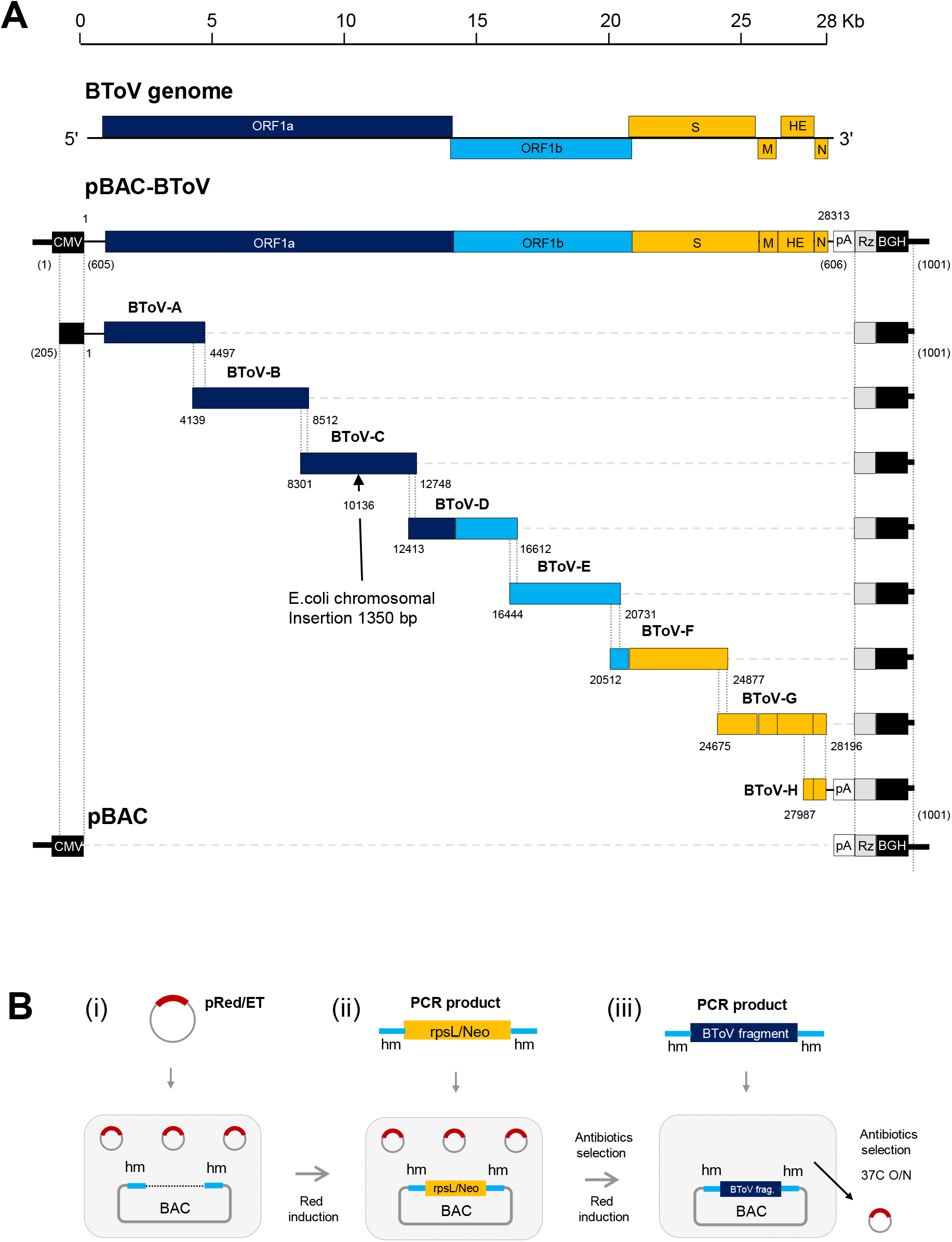
Assembly of a BToV full-length cDNA into a BAC and flowchart of Red/ET recombination. (A) The genome sequence of BToV is divided into eight fragments (BToV-A to H), and each fragment contained homology arms (hm) overlapping each fragment at the 3’ and 5’ terminal ends (vertical dashed line). To distinguish the base number between BToV genome and BAC, the BToV genome of 28,313 nt without poly(A) (25 nt) was written as the normal number, while BAC of 8,219 bp was written as the number in parentheses starting with the CMV promoter. (B) (i) *E. coli* carrying the BAC was electroporated with the pRed/ET plasmid, and the Red/ET recombinant enzyme was induced by L-arabinose. (ii) The linear rpsL/Neo counter-selection/selection cassette flanked by hms was electroporated, and the Red/ET enzyme inserted the rpsL/Neo cassette into the target position via hms. After selecting appropriate antibodies, (iii) the Red/ET enzyme in *E. coli* carrying the modified BAC could replace the rpsL/Neo cassette with the cDNA fragment of BToV in the same way. Only *E. coli* carrying BAC with the target cDNA fragment could be selected by antibodies and 37°C at which the pRed/ET plasmid was removed.

**Fig. 2.**
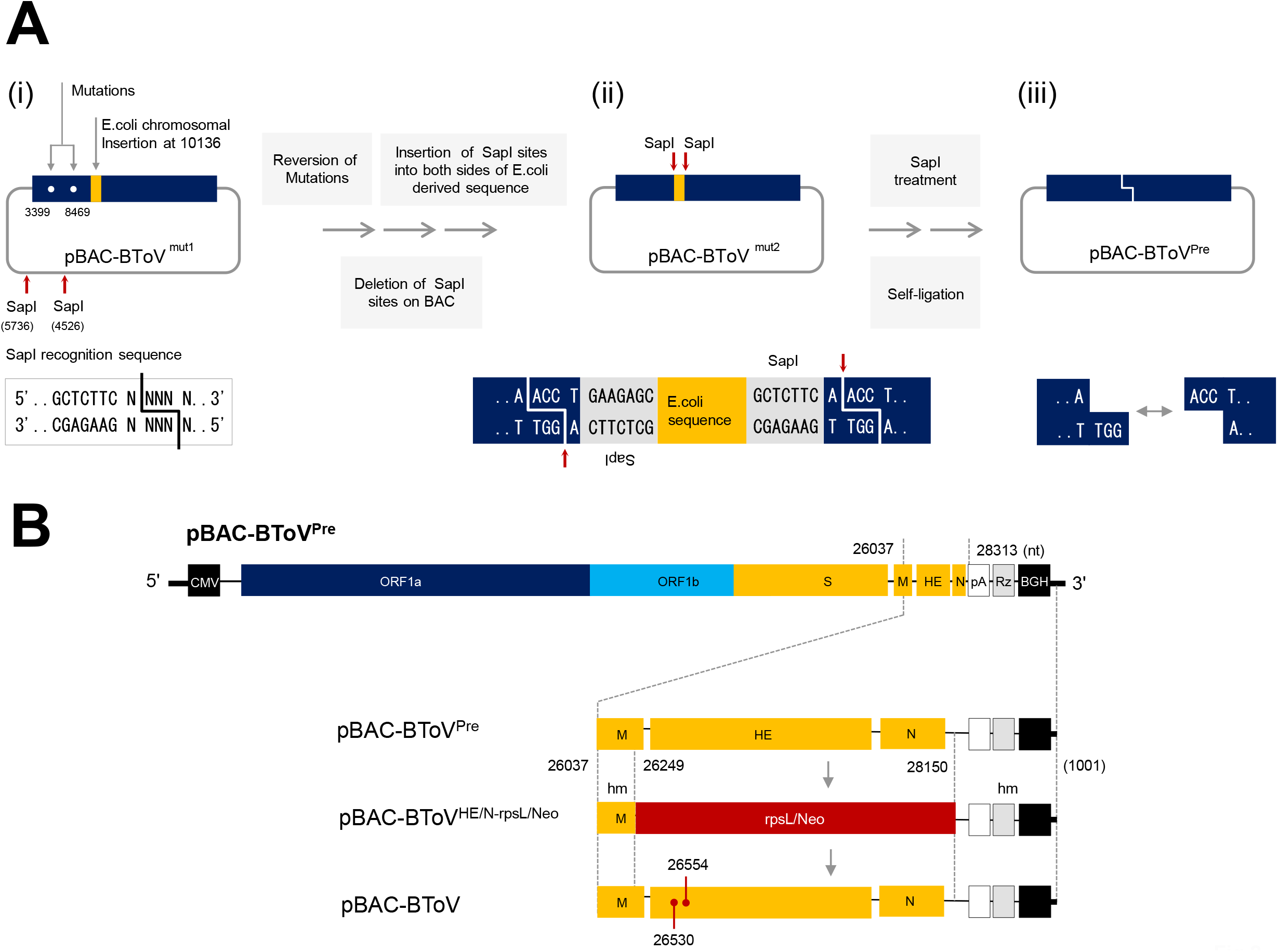
Schematic diagram of pBAC-BToV construction strategy. (A) (i) BAC-BToV^mut1^contained two mutations at nt 3,399 (E→D) and 8,469 (silent), *E. coli* chromosomal insertion at 10,136 and two SapI sites at nt (4,526) and (5,736) on BAC. Two mutations were reverted by the pRed/ET recombination method. To remove the *E. coli*-derived sequence, two SapI sites on BAC were removed by single nt substitution of the recognition site (which is shown below) using in vitro recombination, and (ii) the SapI recognition sites (one is in reverse direction) were inserted on both sides of the E.coli derived sequence by pRed/ET recombination. (iii) Finally, the resulting BAC plasmid (pBAC-BToV^mut2^) was treated with SapI and subjected to in vitro self-ligation (pBAC-BToV^pre^). (B) To introduce genetic makers of pBAC-BToV^pre^, rpsL/Neo cassette was inserted between nt 26,249 and 28,150 containing partial M, HE, and N genes via hms (pBAC-BToV^HE/N-rpsL/Neo^). The rpsL/Neo cassette was replaced with HE carrying the genetic makers at nt 26,530 and 26,554, with the surrounding sequence via the hms (pBAC-BToV).

### Rescue of recombinant BToV

To distinguish between the parent wtBToV and recombinant wtBToV (rBToV) generated from the BAC plasmid, two gene makers were inserted into the HE gene at nt T26530C and T26554A (both silent) (pBAC-BToV) (Fig. 2B). pBAC-BToV was first transfected into nonpermissive 293T, COS7, BHK, Vero, and Huh7 cells because of the very low transfection efficiency of permissive HRT18 cells. Of these cells, 293T showed the best virus production with no clear cytopathic effect (CPE), and the viral titer of the supernatant 3 days post-transfection (dpt) averaged 1.0 × 10^4.1^TCID_50_/ ml (N=3), suggesting that the host range specificity of BToV, which was difficult to propagate in cultured cells, was primarily determined by the entry step, which was also observed in CoVs (47, 59, 63). The supernatant from transfected 293T was inoculated into permissive HRT18 cells, after which the virus was plaque-purified three times, and then the purified virus was used to characterize phonotypic properties. CPE and plaque morphology induced or formed by rBToV were identical to parental wtBToV (Fig. 3B and 4A). Comparison of the full-length genome sequence between rBToV and wtBToV showed complete sequence identities except for two gene markers (Fig. 3C). To compare growth kinetics between rBToV and wtBToV, HRT18 cells were infected with a multiplicity of infection (MOI) of 0.001, and supernatants were harvested at the indicated times. rBToV and wtBToV had indistinguishable growth properties in HRT18 cells and had peaked titers of >1.0 × 10^6.4^ TCID_50_/ml at 48 h post infection (pi) (Fig. 4D). Thus, a full-length infectious cDNA clone of BToV in the BAC system was successfully constructed, and features of the rescued rBToV and parental wtBToV had the same phenotypic properties in HRT18 cells.

**Fig. 3.**
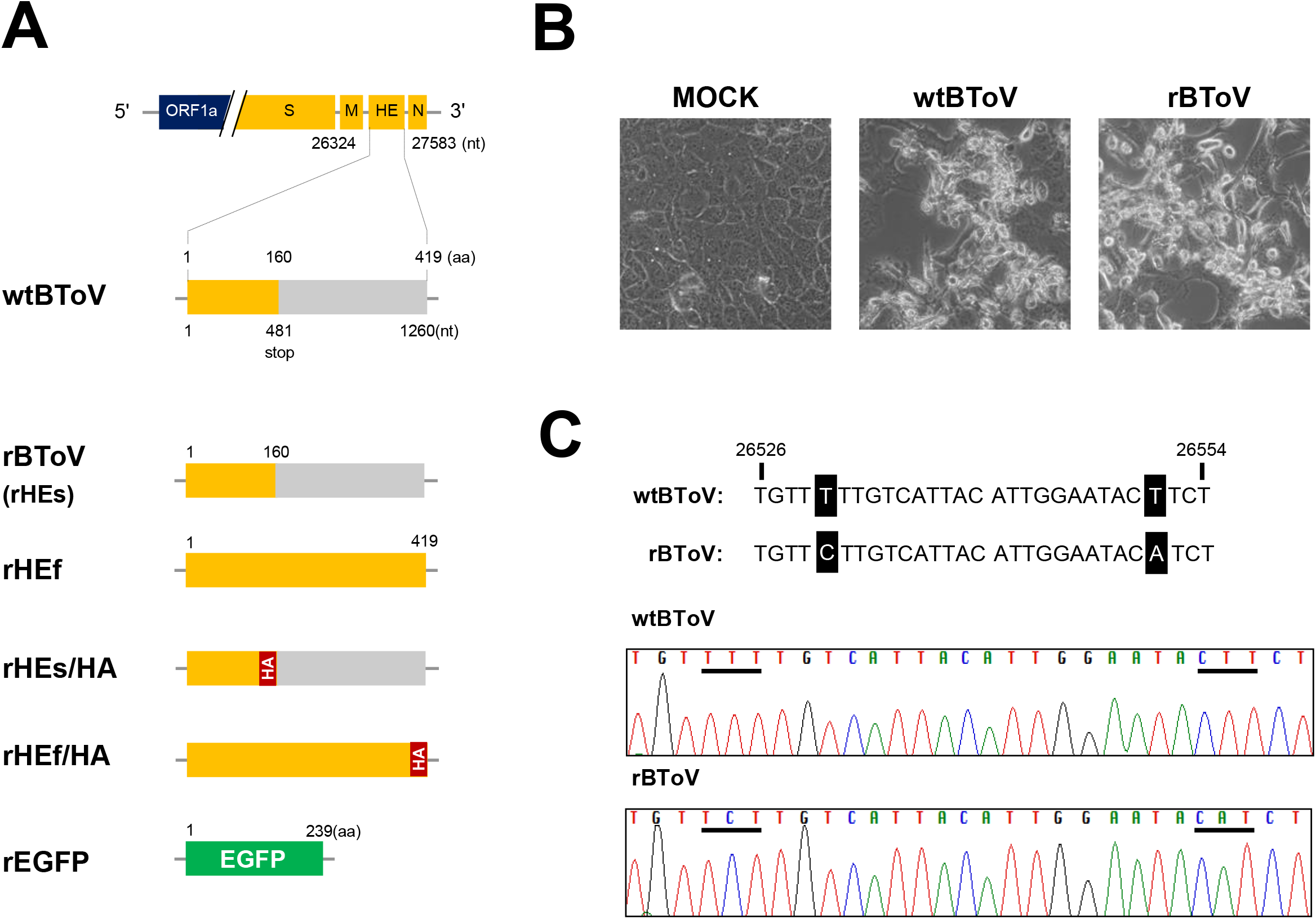
Schematic diagram and generation of recombinant BToVs (A) Cell-adapted wtBToV lost full-length HE (HEf, 419aa) due to a stop codon at nt 481 (the base number of HE gene), resulting in the soluble form HE (HEs, 160aa). rHEf/HA and rHEs/HA were added to the HA-tag at the C-terminal end of HEs and HEf proteins, respectively. rEGFP carries a reporter EGFP gene in which HE gene was completely replaced. nt: nucleotide, aa: amino acid. (B) CPE of HRT18 cells infected with wtBToV and rBToV. HRT18 cells infected with a MOI of 0.05 were observed at 24 hpi under phase-contrast microscopy. (C) Sequence analysis of wtBToV and rBToV at genetic maker sites (underlined) in HE gene.

**Fig. 4.**
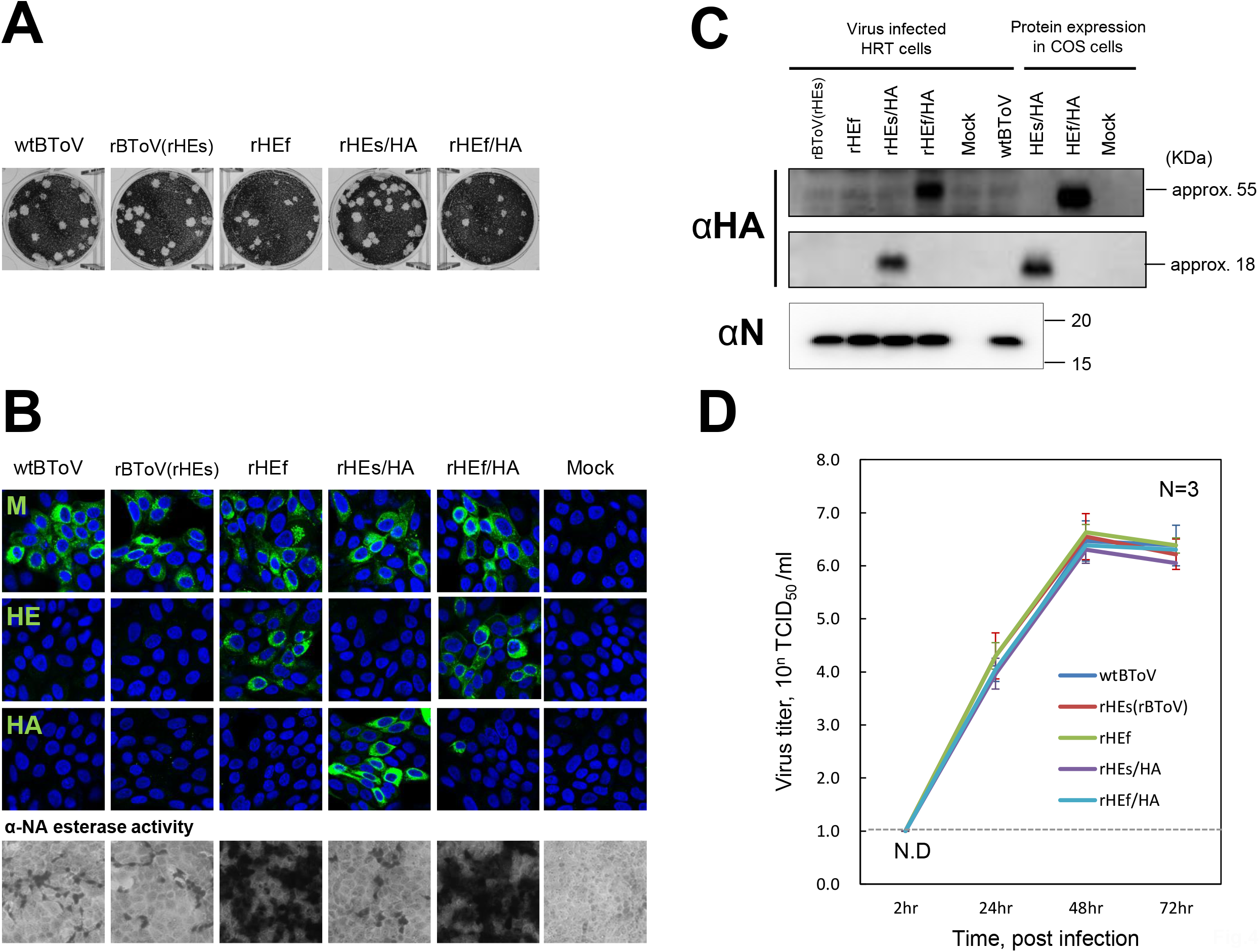
Growth and characteristics of recombinant HE mutant BToVs. (A) Representative of plaque morphology in the 6-well plate of recombinant BToVs at 3 dpi. (B) (Upper) Detection of the membrane (M), HE, and HA-tagged proteins of recombinant BToVs by indirect immunofluorescence. HRT-18 cells were infected with these viruses with a MOI of 0.05 and infected cells were fixed, permeabilized at 24 hpi, and then stained with mouse anti-M and anti-HE antiserum or rabbit anti-HA antibodies and FITC-conjugated secondary antibodies (green). The nuclei were stained with Hoechst (blue). Staining cells were observed under confocal laser scanning microscopy. (Lower) α-NA esterase activity of the cells infected with recombinant BToVs. Fixed infected cells described above were tested for α-NA esterase activity. (C) Immunoblotting of HA-tagged and N proteins in HRT18 cells infected with recombinant BToVs and COS7 cells transfected with plasmid encoding HEs/HA and HEf/HA. Infected HRT18 cells at a MOI of 0.05 at 24 hpi and transfected COS7 cells at 40 hpt were lysed, and the lysates were subjected to immunoblotting. Proteins were detected with rabbit anti-N antiserum and mouse-anti-HA 12CA5 antibody for HEf/HA and rabbit anti-HA antibody for HEs/HA. (D) Growth kinetics of recombinant BToVs. HRT-18 cells were infected with a MOI of 0.001. At the indicated time, supernatants of infected cells were harvested, and virus titers were determined by TCID_50_. The mean and standard deviation of three independent experiments are shown (N = 3). Dashed line shows detection limit. N.D.: not detected.

### Characterization of recombinant BToV with full-length HE or HA-tagged HE gene

Although BToV initially isolated from clinical samples had full-length HE (HEf) genes, cell adapted BToV usually lost HEf due to a stop codon somewhere in HE gene. Cell-adapted wtBToV in our laboratory also has a stop codon at nt 481 (the base number of HE gene), CAG (Q)→<u>T</u>AG (stop), resulting in soluble HE 160 aa in length (HEs) (Fig. 3A). This suggested that HE protein is dispensable for virus replication in cells and may have a negative effect on it (34–37). However, the precise functions of BToV HE protein in cellular viral growth still remains elusive. To investigate HE gene stability and the effect of HE protein on viral growth in HRT18 cells, recombinant BToVs with HEf, or with HA-tagged HEf and HEs genes to facilitate their detection were generated (designated as rHEf, rHEf/HA, rHEs/HA, and rBToV was written as rHEs together here, respectively) (Fig. 3A). All these recombinant BToVs were successfully rescued using the same method described in rBToV. They displayed no significant differences in plaque morphology and CPEs (data not shown) in HRT18 cells compared to parental wtBToV and rBToV (rHEs) (Fig. 4A). To detect HEf or HA-tagged HEf and HEs proteins, HRT18 cells infected with recombinant BToVs were fixed at 24 hpi. The cells were then subjected to an α-NA esterase assay or to indirect immunofluorescence (IF) using mouse anti-M and anti-HE antiserum or using rabbit anti-HA and mouse anti-HA 12CA5 antibodies. Stained cells were observed under confocal laser scanning microscopy (Fig. 4B). M proteins were detected similarly in all cells infected with the recombinant BToVs, and HE proteins and α-NA esterase activity were observed only in cells infected with rHEf and rHEf/HA. On the other hand, HEs/HA proteins were well stained by both anti-HA antibodies (only rabbit anti-HA antibody was shown), whereas HEf/HA protein was barely stained despite being well detected by mouse anti-HE antiserum (Fig. 4B). We next tried to detect HEf/HA by immunoblotting. Infected cells were lysed, and detection was performed using rabbit anti-N antiserum and two anti-HA antibodies. With similar detection of N proteins in all infected cells, HEs/HA could be detected in both anti-HA antibodies similar to IF results, whereas HEf/HA could not be detected by rabbit anti-HA but could only be detected by the mouse anti-HA 12CA5 antibody (Fig. 4C). To investigate whether the different reactivity of the C-terminal HA-tag was due to the modification of the HA-tag in the cytoplasm by either an infected cell metabolic change or viral proteins, HA-tagged proteins in COS7 cells transiently expressing HEf/HA and HEs/HA were attempted to be detected in the same way. Both HA-tagged proteins could be stained in IF (data not shown), but reactivity did not change in immunoblotting (Fig. 4C). Although we could not rule out the possibility that differences were due to the modification or due to the expression level of HA-tag of HEf/HA in infected cells from IF results, immunoblotting results at least suggested that the C-terminal HA-tag of HEf/HA had the specific structure that affects reactivity of the anti-HA antibodies. Finally, to compare growth kinetics, HRT18 cells were infected with these recombinant BToVs with a MOI of 0.001. All these recombinant BToVs, wtBToV, and rBToV (rHEs) had no significant differences in growth properties (Fig. 4D). These results indicated that recombinant BToVs could express full-length HE and HA-tagged proteins in infected cells with growth ability comparable to those of the parental wtBToV and to rBToV (rHEs)

### Full-length HE gene stability of rHEf and rHEf/HA

To assess stability of the full-length HE gene of rHEf and rHEf/HA, HRT18 cells were infected with two clones (No.1 and No.2) derived from different plaques of each virus, and viruses were serially passaged until twenty passages (P20). Cells were infected with the viruses harvested at the indicated passage history and were fixed or lysed at 24 hpi, and then subjected to either α-NA esterase assay or N protein-detection by immunoblotting (Fig. 5A and 5B). N proteins were detected in all infected cells (Fig. 5B). On the other hand, cells infected with rHEf No.2 and two rHEf/HA retained α-NA esterase activity until P19, whereas cells infected with rHEf No.1 gradually decreased in activity during passages and completely lost activity at P19 (Fig. 5A). RT-PCR of the supernatants from infected cells (resulting in +1 passage history) using primers flanking the HE gene showed that during serial passages, band sizes of rHEf No.1 without esterase activity and rHEf No.2 with esterase activity were not different (Fig. 5C). On the other hand, although both rHEf/HA exhibited esterase activity, a smaller band was observed in rHEf/HA No.2 from P10. To further study their HE genes, all these viruses at P19 were plaque-purified once, and the HE gene of viruses from six well-isolated plaques were sequenced. As summarized in Fig. 5D, all six rHEf No.1 sequences had one base deletion at nt T19 in the HE gene, causing 17 aa short peptide with a stop codon by -1 frame shifting, and three out of six had an additional deletion between nt 1,216 to nt 1,241. In contrast, all six rHEf No.2 retained the full-length HE gene, and one out of six had an amino acid substitution at T321I in the HE protein. Out of the six, five rHEf/HA No.1 retained the HA-tagged full-length HE gene, but one had an aa mutation within the HA-tag (YPYDYPDYA→Y<u>L</u>YDYPDYA), whereas three rHEf/HA No.2 retained the HA-tagged full-length HE gene with a D247E substitution in the HE protein in one virus but the other three had a deletion between nt 278 to nt 878, causing a short soluble HE protein 99 aa in length. These data partly supported the idea that HE proteins are not essential or could have a negative effect on virus growth in HRT18 cells. However, it is notable that each one clone could retain the full-length HE gene up to at least twenty passages. Thus, cell adapted BToV may be able to retain full-length HE gene more stably in cells under certain conditions.

**Fig. 5.**
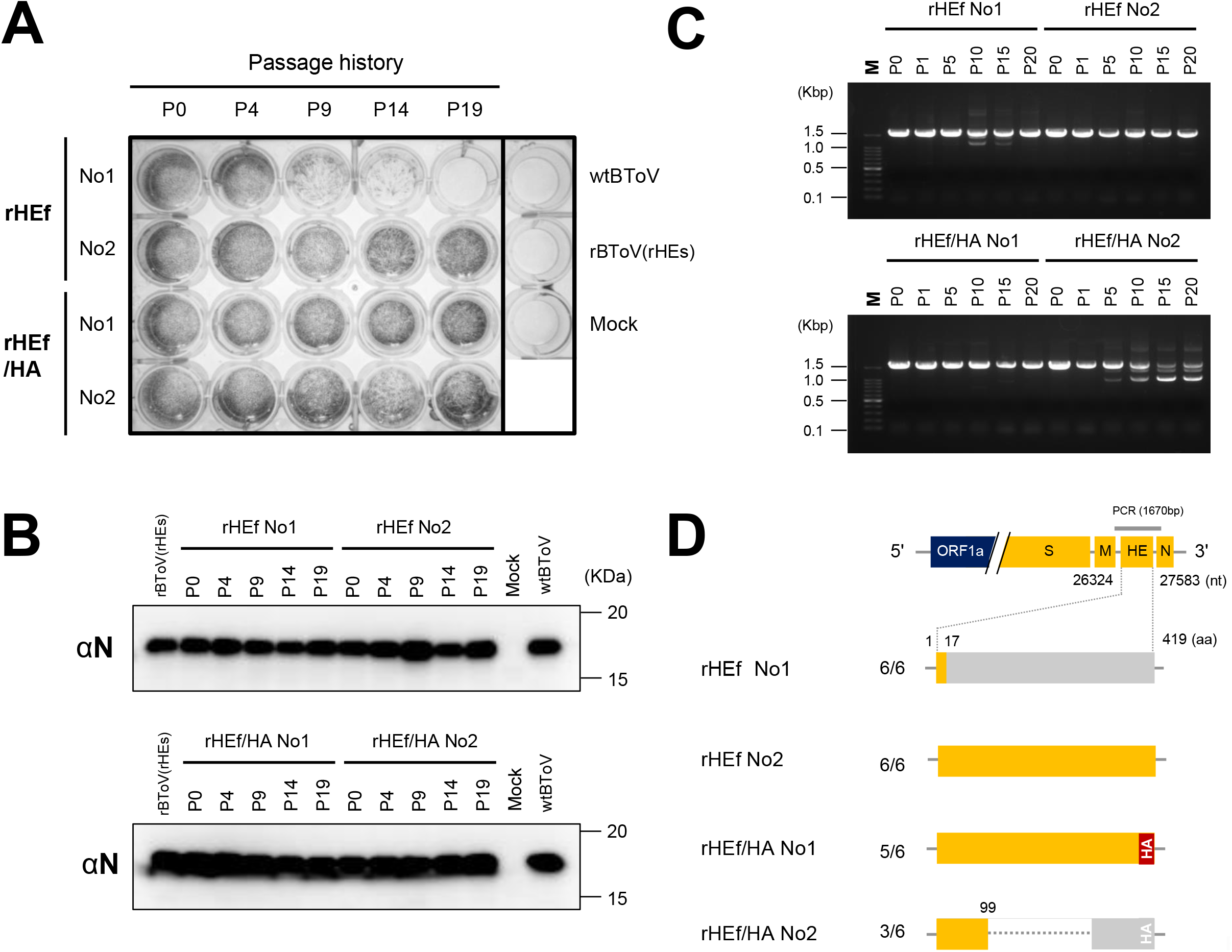
Full-length HE gene stability of recombinant HE mutant BToVs. (A) α-NA esterase activity of infected cells with recombinant BToVs carrying full-length HE gene during serial passages. HRT-18 cells in 24-well plate infected with a MOI of 0.05 were fixed at 24 hpi, and then tested for α-NA esterase activity. Two clones (No.1 and No.2) derived from different plaques of each virus were used. (B) Immunoblotting of N proteins in HRT18 cells infected with recombinant BToVs described above at the indicated passage history. (C) RT-PCR of the HE gene of recombinant BToVs. The supernatant from infected cells (resulting in +1 passage history) were amplified by RT-PCR using primers flanking the HE gene (shown in figure D, 1670 bp). Amplicons were analyzed by agarose gel electrophoresis. (D) Sequencing analysis results summary of recombinant BToVs HE genes. Viruses at P19 were plaque-purified once and six well-isolated plaques were picked, and then these viruses were inoculated into fresh HRT18 cells. The HE gene of each supernatant was amplified and sequenced. 5/6 means that 5 out of 6 plaques have the HE proteins described in figure.

### Characterization of recombinant BToV carrying EGFP gene

Since recombinant viruses carrying the reporter gene are helpful in understanding fundamental viral processes and for screening therapeutic compounds, we attempted to generate recombinant BToV by replacing the HE gene with the EGFP gene (rEGFP). Rescued rEGFP were plaque-purified three times and expanded (clone No.1 and No.2) or expanded without plaque purification (No.3 to No.6) because of the instability of the EGFP gene described below. rEGFP formed smaller and more heterogeneous plaques compared to the parent wtBToV (Fig. 6A). To detect EGFP in infected cells, infected cells at 36 hpi were observed under fluorescence microscope, or fixed, permeabilized, stained with mouse anti-M antiserum, and observed under confocal laser scanning microscopy (Fig. 6B). EGFP-expressions were observed only in cells infected with rEGFP but not with wtBToV and rBToV, and cells expressing EGFP always also expressed M protein (Fig. 6B). These infected cells were next subjected to N protein and EGFP detection by immunoblotting. Although the amount of N protein in the rEGFP-infected cells was significantly less than that of cells infected with wtBToV and rBToV, EGFP was only detected in rEGFP-infected cells (Fig. 6C). Comparison of growth kinetics among rEGFP, wtBToV, and rBToV showed that rEGFP significantly decreased growth ability. In four independent experiments, no virus could be detected at 24 hpi, and peak titer 1.0 × 10^3.8-4.6^ TCID_50_/ml was observed at 48 to 72 hpi in three experiments, which decreased by >2 log compared to parent viruses (Fig. 6D). In one experiment, peak titer reached 1.0 × 10^6.6^ TCID_50_/ml at 72 hpi, but the virus no longer expressed EGFP (Fig. 6D). These data indicated that rEGFP could express EGFP in infected cells but significantly decreased its growth properties. Moreover, rEGFP seemed to readily lose EGFP-expression during virus growth.

**Fig. 6.**
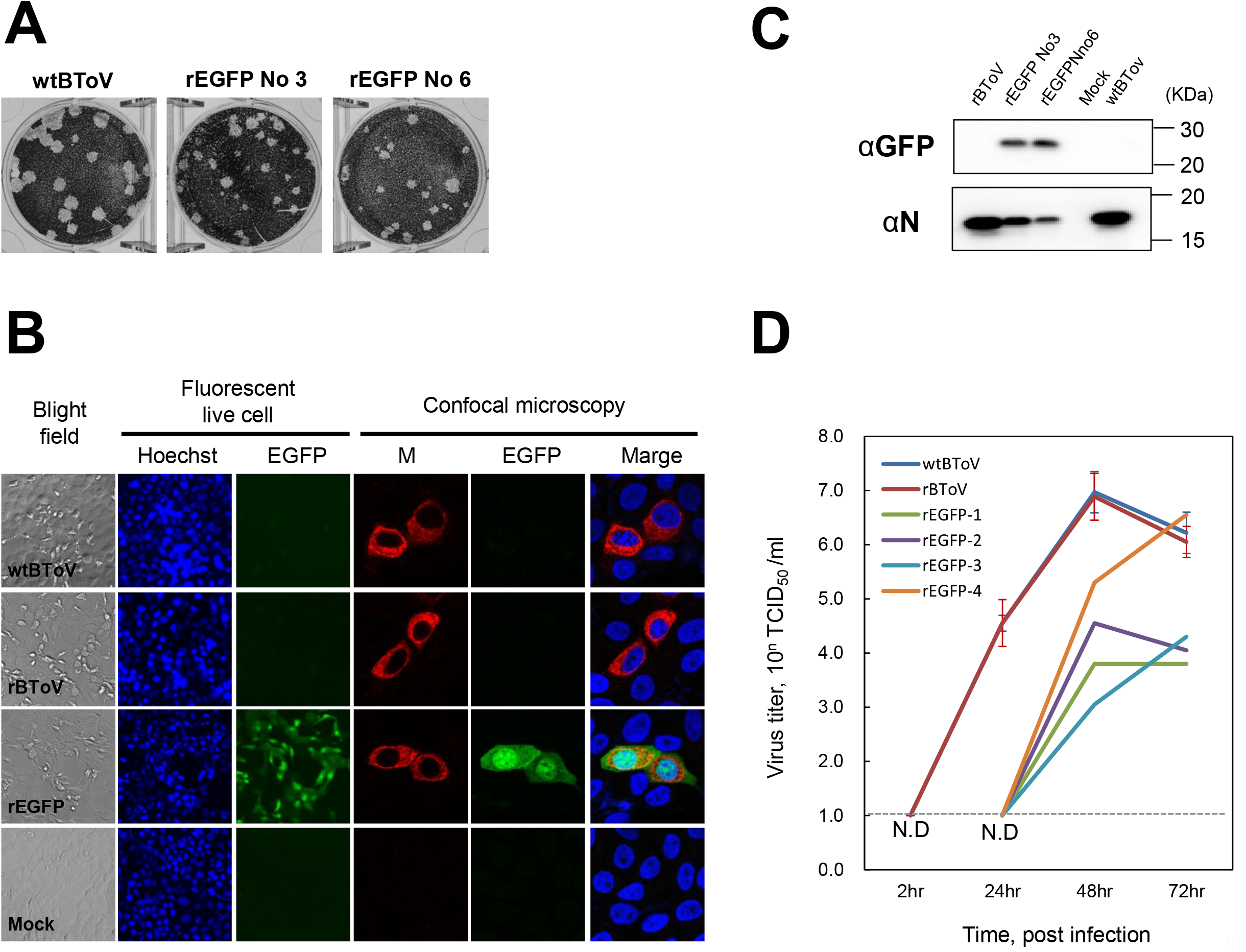
Growth and characteristics of recombinant BToV carrying EGFP gene. (A) Representative of plaque morphology of rEGFP at 3 dpi. In these rEGFP, 293T cells in 3 cm dish were transfected with BAC-BToV-EGFP, and the supernatants harvested after 2 dpt were inoculated into fresh HRT18 cells in a 10 cm dish without plaque purification. This was stored as passage 0 (P0) virus, and the viruses obtained from each independent experiment were numbered (No.3 to No.6) (B) Detection of EGFP-expression (green) in infected cells with rEGFP under a fluorescence microscope. HRT18 cells were infected with rEGFP No.3 with a MOI of 0.05, and infected cells were stained with Hoechst (Blue) at 36 hpi. Living infected cells were observed under a fluorescence microscope (left). Infected cells were fixed at 36 hpi, permeabilized, stained with mouse anti-M antiserum and AlexaFluor™ 594-conjugated secondary antibody (red), and observed by confocal laser scanning microscopy (right). Cells infected with rBToV, wtBToV were observed or fixed at 24 hpi. (C) Immunoblotting of EGFP and N proteins in infected cells with rEGFP. Infected cells described above were lysed at 36 hpi. Cells infected with rBToV and wtBToV were lysed at 24 hpi. Proteins in the lysates were detected with rabbit anti-N antiserum and mouse anti-GFP antibody. (D) Growth kinetics of rEGFP No.3. Virus titers were determined as described in Fig. 4. Each result of four independent experiments of rEGFP No.3 is shown. Dashed line shows detection limit. N.D.: not detected.

To study EGFP gene stability, HRT18 cells were infected with rEGFP, and viruses were serially passaged until five passages (P5). Cells were infected with rEGFP harvested at the indicated passage history and were observed under a fluorescence microscope (Fig. 7A). As a result, EGFP was not observed in most of the cells infected with rEGFP (No.1 to No.4) after one passage. RT-PCR of P0 and P1 viruses using primers flanking EGFP gene found that the all P0 rEGFP predominantly had the full-length EGFP gene, but smaller bands were observed in all P1 rEGFP (No.1 to No.4) (Fig. 7B). Of these, the major bands of P1 rEGFP (No.2 to No.4) were analyzed, and it was found that a large part of the EGFP gene was deleted (Fig. 7D), suggesting that rEGFP readily lost EGFP-expression due to deletion of EGFP gene after one passage.

**Fig. 7.**
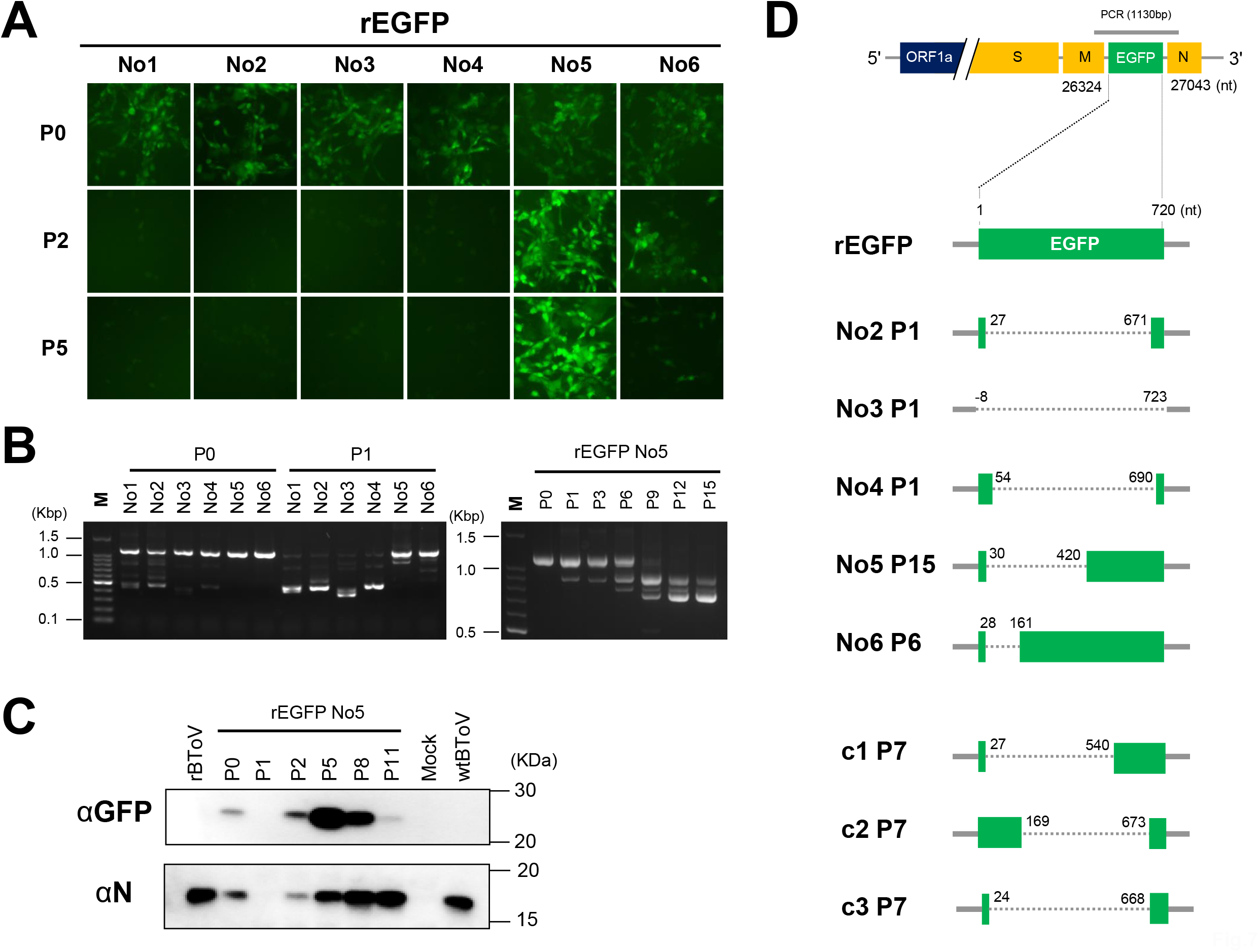
EGFP gene stability of rEGFP. (A) EGFP-expression of rEGFP during serial passages. HRT-18 cells were infected with a MOI of 0.001, and then 1,000-fold diluted virus was inoculated into fresh cells. Viruses were then harvested after 3 dpi. Two rEGFP (No.1 and No.2) were plaque-purified and cloned, while four rEGFP (No.3 to No.6) described in Fig. 6 were not. Cells were infected with 1,000-fold diluted rEGFP harvested at the indicated passage history and observed at 36 hpi under a fluorescence microscope. In P0 viruses, infected cells with a MOI of 0.05 (actual experiment was 0.001) are shown to easily observe EGFP-expression. (B) RT-PCR of the EGFP gene of rEGFP. The supernatant viruses from infected cells (+1 passage history) were amplified by RT-PCR using primers flanking the EGFP gene (Figure D, 1130bp). Amplicons were analyzed by agarose gel electrophoresis. (C) Immunoblotting of EGFP and N proteins in cells infected with rEGFP No.5 at indicated passage history. Cells were infected with 1,000-fold diluted rEGFP No.5 from P1 and infected with P0 viruses (rEGFP No5, rBToV, wtBToV) at a MOI of 0.05. Infected cells with rBToV and wtBToV and with rEGFP No.5 were lysed at 24 hpi and at 36 hpi, respectively. (D) Summary of EGFP gene sequencing analysis of rEGFP. Major bands of EGFP gene of each supernatant were purified and sequenced.

### Improvement of EGFP-expression and growth properties in rEGFP variant

Interestingly, rEGFP No.5 and No.6 maintained the EGFP gene after one passage. No.6 gradually decreased EGFP-expression until P5, while No.5 appeared to gradually increase (Fig. 7A). Thus, serial passages of rEGFP No.5 was continued until P15, and the viruses were then harvested. Cells were infected with rEGFP No.5 harvested at the indicated passage history and their supernatants (+1 passage history) were collected and subjected to RT-PCR targeting EGFP gene (Fig. 7B), or the infected cells were lysed at 36 hpi and subjected to N protein and EGFP detection by immunoblotting (Fig.7C). Although smaller bands were observed, rEGFP No.5 retained full-length EGFP gene at least until P6 and mainly lost it by P9 (Fig. 7B), but EGFP-expression was still observed in infected cells even at P9 to an extent (data not shown). The major bands of P15 rEGFP No.5 (and P6 No.6: data not shown) were analyzed by sequencing, and a part of the EGFP gene was deleted (Fig. 7D). Similar to RT-PCR results, immunoblotting of N protein and EGFP of rEGFP showed that EGFP-expression level of No.5 was maximized at P5, and this gradually decreased until P11 (Fig. 7C). It should be noted that the EGFP-expression level or growth property seemed to dramatically increase compared to that of P0 rEGFP No.5.

To further investigate P5 rEGFP No.5, this was plaque-purified three times and three clones were expanded (designated rEGFP c1-c3). These variants formed similar plaque morphologies to wtBToV and rBToV (Fig. 8A). Comparing growth kinetics among rEGFP c1-c3, wtBToV, and rBToV showed that their peak titer reached 1.0 × 10^6.5-7.2^ TCID_50_/ ml at 72 hpi, which is comparable to that of the parental viruses, although the peak was reached one day slower (Fig. 8B). Cells infected with rEGFP c1-c3 were lysed at 24 hpi, and these lysates were then subjected to N protein and EGFP detection. The result indicated that EGFP-expression levels in these rEGFP c1-c3 were dramatically improved compared to P0 original virus (Fig.8C). Analysis of the full-length genome sequence of rEGFP c1 revealed five aa substitutions at G1001S, C1442F, T2129I in NSP1, I3562T in NSP4, and I5327V in NSP10 (Helicase), respectively, strongly suggesting that these accumulated mutations in NSPs were involved in this phenotype (Fig.8D). Finally, EGFP gene stability in rEGFP c1-c3 were investigated. As shown in Fig. 8E, these variants tend to stably express EGFP until P4, but smaller bands due to the deletion of the full-length HE gene were mainly observed, with EGFP-expression decreasing until P6 (data not shown in rEGFP c3). Sequencing the major band of P7 rEGFP c1-c3 showed deletion of the EGFP gene, similar to those observed in other rEGFP (Fig.7D). These data indicated that any or all of the five mutations in NSPs would contribute to some acceptance of exogenous gene such as EGFP gene, resulting in improved EGFP-expression and viral growth in cells, although the EGFP gene was eventually lost

**Fig. 8.**
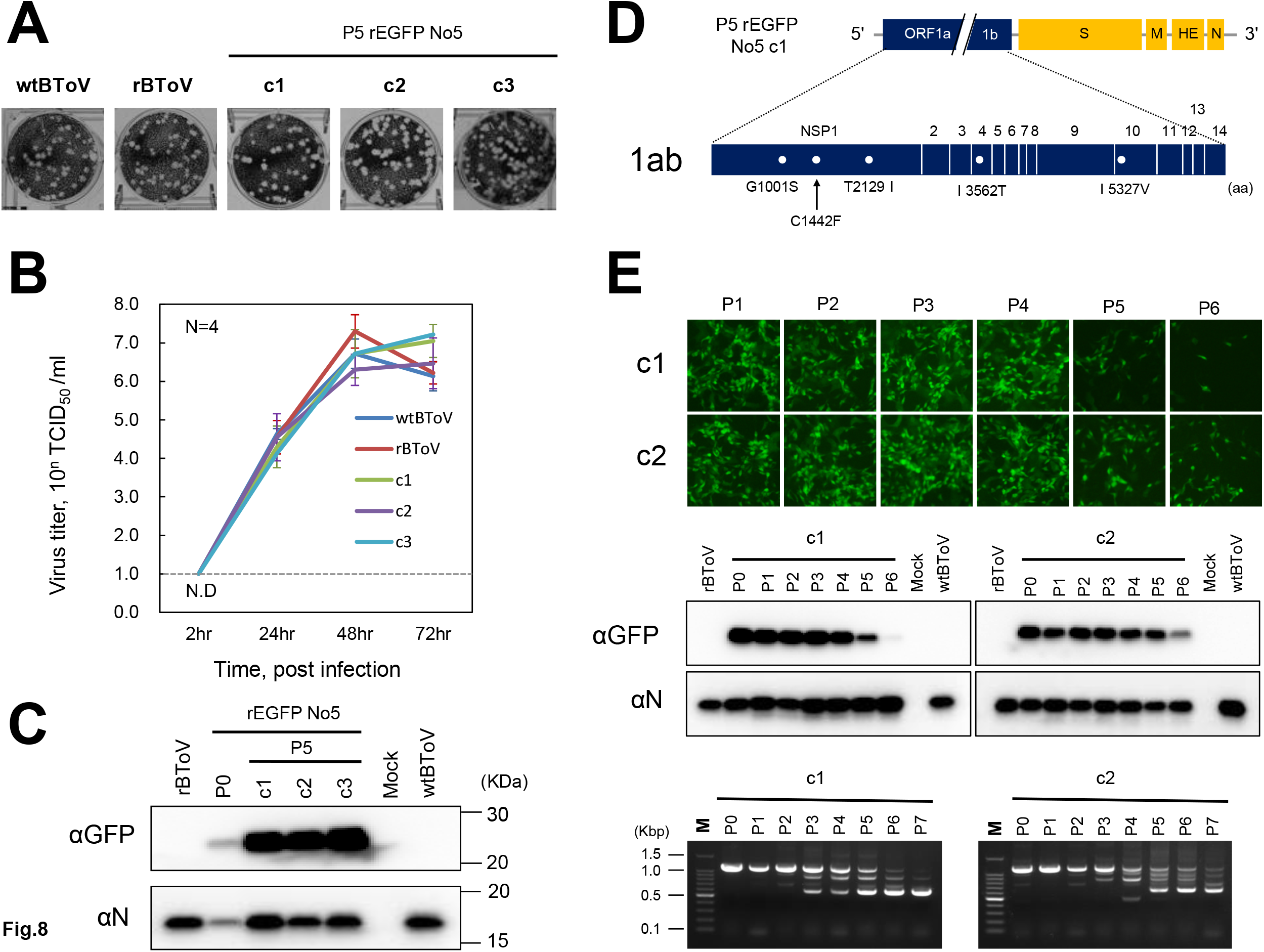
Improvement of the growth kinetics and EGFP-expression of rEGFP. (A) Representative of plaque morphology of variant rEGFP. The P5 rEGFP No.5 was plaque-purified three times, and three clones were expanded (designated c1 to c3). (B, C) Growth kinetics and immunoblotting of rEGFP (c1 to c3). Virus titers (N=4) were determined and immunoblotting was performed as described in Fig. 6. Dash line shows detection limit. N.D.: not detected. The cells were infected with a MOI of 0.05, and lysed at 24 hpi for variant rEGFP rBToV, wtBToV and at 36 hpi for P0 rEGFP No.5. (D) Full-length genome sequence of rEGFP c1. Mutations different from the pBAC-BToV-EGFP are shown. (E) Genetic stability of EGFP gene of rEGFP c1 and c2. Analysis of EGFP-expression and EGFP gene of these viruses during serial passages are described in Fig.7

## Discussion

In this study, we successfully constructed a full-length infectious BToV cDNA clone in a bacterial artificial chromosome. To our knowledge, this is the first reverse genetics system for the new family *Tobaniviridae*.

### BAC construction

Red/ET recombination method used to assemble eight BToV cDNA fragments into the BAC has been recently utilized for full-length cDNA assembly of some CoVs, PEDV, and FIPV (58, 75) and for introducing mutations (76–78). This method is more rapid and produces mutated clones more efficiently than traditional methods (78); however, despite several tries, the unintentional insertion of an *E. coli*-derived sequence which may counter toxic regions could not be removed. Red/ET seemed not to function in restoring the toxic region to the host. Thus, a type IIS restriction enzyme, SapI, was inserted into both ends of the *E. coli*-derived sequence, and this was removed by in vitro self-ligation. The resulting plasmid could be successfully propagated in *E. coli* without pRed/ET enzyme. Similar observations were not seen in the full-length genome assembly of PEDV and FIPV into low copy BAC plasmid (58, 75), but were seen in the subcloning of the TGEV cDNA fragment into high copy plasmids (47). The position wherein the *E. coli* sequence is inserted is similar in both (BToV at nt 10,136 and TGEV at nt 9,973), suggesting that these regions (BToV is in 3C-like protease) were particularly toxic in bacteria in ToVs and CoVs. Though it remains unclear whether very small amounts of 3C-like protease in *E. coli* or, functional viral proteins cleaved by this protease causes toxicity, the *E. coli*-derived sequence was inserted not when the BToV-C fragment containing the 3C-like protease gene was assembled but when the full-length 1ab gene (BToV-F fragment) was assembled. For BToV, since insertion of the *E. coli*-derived sequence was observed even in low copy BAC plasmid, this region may be more toxic than that of CoV or Red/ET recombination method may be prone to this. Thus, to establish other BAC-based ToV reverse genetics using this method, it may be better to counter the toxic region (3-C like protease gene) beforehand with an *E. coli*-derived sequence flanked by type IIS restriction enzymes for subsequent in vitro ligation.

### Characterization of recombinant BToV with manipulating HE gene

Since cell-adapted BToV generally lost the full-length HE gene, HE protein is not essential for viral growth in cells or could suppress growth (34–37, 69). To study this, we generated rHEf with full-length HE gene and rHEf/HA, rHEs/HA with HA-tagged HEf, or HEs gene. Comparing their growth kinetics to wtBToV and rBToV (rHEs) showed no significant differences; however, during serial passaging of two clones from each rHEf and rHEf/HA, each clone lost the full-length HE gene, supporting the above idea. In contrast, each the other clone could retain the full-length-HE gene at least until twenty passages. This was unexpected because another BToV (Niigata strain) isolated from a clinical sample in our laboratory easily lost the full-length HE gene somewhere due to a stop codon insertion until eight passages during cell-adaptation (Islam and Taguchi, unpublished data). Thus, the negative effect on HE protein in the cells of cell-adapted Aichi-BToV may be lower than that of non-cell-adapted Niigata-BToV. Furthermore, HE proteins of cell-adapted Aichi-BToV may have some advantage in terms of viral growth in the cells.

HE proteins are retained not only in ToV, but also in some of the beta CoVs (79). HE is a type I glycoprotein on the viral envelope with two reversible functional domains: one binds O-acetylated sialic acids (O-Ac-Sia) and the other destroys this binding. Both viruses also contain another type I glycoprotein S protein responsible for receptor-binding and fusion activity. Among some beta CoVs, the species *Betacoronavirus-1*, including human coronavirus OC43 (HCoV-OC43), bovine coronavirus, and porcine hemagglutinating encephalomyelitis virus, use O-Ac-Sia as the receptor via the S protein (80, 81). Thus, these viruses bind to O-Ac-Sia via both the S and HE proteins, while HE acts as a receptor-destroying enzyme that promotes release of progeny viruses from infected cells and helps avoid attachment to non-permissive cells or to receptor-like decoy. The importance of HE in these viruses has been reported, and HE of HCoV-OC43 plays an essential role in the efficient release of progeny viruses from infected cells (82) and balancing receptor-binding and receptor-destroying of HE contributes to host adaptation (83). In contrast, the species *Murine coronavirus*, a mouse hepatitis virus (MHV) in which the S protein uses CAECAM1a as receptor (84) and in which HE binds to O-Ac-Sia exclusively (85) does not require HE proteins for viral growth. In fact, many cell-adapted laboratory strains of MHV fail to produce HE (86), with MHV rapidly losing it during serial passages (87, 88), which is similar to BToV. On the other hand, experiments using recombinant MHV carrying the full-length HE gene showed about half of the HE deficiencies until six passages (88), while our experiment showed that some recombinant BToVs could retain HE at least until twenty passages. This may be explained by the hemagglutination activity of the S protein of cell-adapted Aichi-BToV (37). It is not known whether the hemagglutination activity is mediated via O-Ac-Sia binding, but, if so, HE protein may contribute to the regulation of S protein-mediated cell surface attachment to facilitate release of progeny viruses. Alternatively, in twenty passages, accumulation of mutations in other regions may contribute to HE protein acceptance, as observed in rEGFP. Although the precise function of BToV HE remains to be elucidated, further studies using these recombinant BToVs will bring better understanding.

### Characterization of recombinant BToV with EGFP gene

Since recombinant viruses expressing reporter proteins such as GFP, Red Fluorescent Protein and luciferase are useful for understanding fundamental viral processes and for high-throughput testing of therapeutic compounds, some recombinant CoVs carrying these reporters were generated (50, 54, 55, 57, 58). These recombinant CoVs have their ORF3, ORF4 or ORF5 accessory genes replaced with reporter genes, and these could all stably express reporter proteins, with their growth abilities comparable to or slightly reduced compared to the parent virus. In contrast, rFGFP in which the HE gene, which is also an accessory gene, was substituted with EGFP gene, could express a fluorescent protein, but it showed significantly reduced growth compared to parental wtBToV, as well as less EGFP-expression. Furthermore, rEGFP easily lost EGFP-expression ability after only one passage due to the deletion of HE genes. Since EGFP itself is a neutral protein, which probably does not give any advantage or disadvantage to viral growth in cells, whereas rEGFP with deleted EGFP gene increased in terms of growth properties (Fig.6D), EGFP gene itself rather than EGFP would lead to a significant decrease in viral growth. It is interesting to note that, despite eight independent experiments, starting points of HE deletion in the five were concentrated at around specific positions, namely nt 24, 27, 28, and 30, and likewise the end points of the four were concentrated at around nt 668, 671, 673, and 690 (Fig.7D), suggesting that “specific negative positions” of the EGFP gene are easily recognized and deleted. Deletion patterns of the EGFP gene showed that in many cases, a large part of or the entire EGFP, even including a part of the transcription*-*regulating sequence of EGFP (HE) and, the untranslated region between EGFP (HE) and the N genes, were deleted, whereas a relatively short deletion leading to retention of more than 80% of the EGFP gene was also observed (Fig. 7D). All rEGFP with deleted genes rapidly replaced intact rEGFP, likely under negative selection of EGFP genetic structure. These suggested that either the entirety or a part of the EGFP gene structure may suppress viral genome replication or transcription, resulting in decreased viral growth.

On the other hand, rEGFP variant obtained during serial passages showed remarkably improved EGFP-expression and growth ability in the cells. This is noteworthy considering EGFP gene has a negative effect on viral growth whereas a neutral EGFP protein exists without clear positive selection. rEGFP variant contained accumulated five mutations in NSPs: three in NSP1, and one each in NSP4 and NSP10 (helicase), respectively. Though functions of NSP1 and NSP4 of ToVs remain unknown, the primary function of helicase, which is highly conserved among nidoviruses including ToVs and CoVs, is to unwind DNA or RNA duplexes (89). The crystal structure of MERS-CoV helicase (corresponding to NSP13) showed that it comprises multiple domains, an N-terminal Cys/His rich domain (CH) binding to three zinc atoms, a beta-barrel domain, and a C-terminal helicase core with two Rec-A like domains (90). Protein sequence alignment between MERS and BToV helicase by MEGA6 software (91) revealed that the mutation at I5327V (I68V in helicase) was located next to Cys69 (corresponding to Cys72 of MERS-CoV helicase) in the CH, one of the cysteine residues that coordinated with the third zinc atom. It was reported that alanine substitutions in highly conserved Cys/His in CH of HCoV-229E and other nidovirus equine viral arteritis have big effects on helicase activity (92, 93). Thus, one possibility is that mutation in the CH of rEGFP helicase may facilitate to unwind “specific negative positions” of the EGFP gene, resulting in increased growth. However, even this variant showed similar EGFP gene deletion patterns to other intact rFGFP until five passages. This finding that the virus tolerates unfavorable foreign gene to some degree without clear positive selection may contribute in understanding the mechanism by which the virus evolves by accepting other genes.

In this article, we descraibe a reverse genetics platform for BAC-based BToV, which is a useful tool for understanding fundamental viral processes and pathogenesis and for BToV vaccine development. Moreover, since the rEGFP variant carrying the middle stable EGFP gene can be plaque-purified and expanded for practical use, this recombinant virus can be used for high-throughput testing of therapeutic compounds and entry assays. Recently, in addition to a unique combination of discontinuous and continuous ToV RNA synthesis(94), differences between CoVs and ToVs such as “hidden” proteins (U1 and U2) of unknown function suspected to be translated from non-AUG initiation codons (CUG-initiation) (95) and N protein localization, have been highlighted. This reverse genetics system can provide future understanding of ToV proteins and unique RNA synthesis mechanisms.

## Materials and Methods

### Cells and Viruses

Human rectal adenocarcinoma subcell line (HRT18-Aichi) (34, 36, 69), 293T, and COS7 cells were maintained at 37°C and 5% CO_2_ in Dulbecco’s Modified Minimum Eagle Medium (DMEM) (Sigma-Aldrich, MO, USA) supplemented with 10% fetal bovine serum (FBS) [DMEM(+)] and penicillin (50 units/mL)-streptomycin (50 µg/mL) (PS) (Sigma-Aldrich). Cell-adapted BToV (Aichi strain) was propagated in HRT18 cells in DMEM without FBS [DMEM(-)], as FBS inhibits BToV infection.

### Antibodies

Anti-HE, anti-M polyclonal mouse (mouse anti-HE and anti-M) antiserum, and anti-N polyclonal rabbit (rabbit anti-N) antiserum were obtained by in vivo electroporation of mice with pCAGGS-BToV-HEf according to the method previously described (70), by immunizing mice with the M protein synthetic peptide (2-FETNYWPFPDQAPN-15), and by immunizing rabbit with the N protein synthetic peptide (141-EVSSGDQETPHKIA-154). Anti-HA-tag mouse monoclonal antibody (Clone: 12CA5) (mouse anti-HA 12CA5) was obtained from supernatants of the hybridoma cells. Anti-HA-tag polyclonal rabbit IgG (rabbit anti-HA) antibody (MBL Nagoya, Japan) and anti-GFP mouse monoclonal antibody (mFx75) (mouse anti-GFP) (Fujifilm Wako, Osaka, Japan) were purchased. Secondary antibodies, horseradish peroxidase (HRP)-conjugated goat anti-rabbit IgG (Thermo Fisher Scientific, MA, USA), HRP-conjugated goat anti-mouse IgG (Rockland Immunochemicals, PA, USA), Alexa Fluor™ 594-conjugated goat anti-mouse IgM and IgG (Thermo Fisher Scientific), FITC-conjugated goat anti-rabbit IgG (MP Biomedicals, CA, USA), and FITC-conjugated goat anti-mouse IgG+IgM+IgA (Rockland Immunochemicals) and IgG+IgM (Jackson Immuno Research, PA, USA) were purchased. Two Alexa Fluor™ 594-conjugated goat anti-mouse and two FITC-conjugated goat anti-mouse antibodies were mixed 1:1, respectively, and were used.

### Plaque assay

HRT18 cells were seeded onto 6-well plates with 1.0 × 10^6^ cells/well. The next day, the cells were inoculated with 400 μL of diluted virus in order to form 10–20 plaques per well and incubated at 37°C for 1.5 h. Unabsorbed virus were removed and cells were overlaid with MEM (Sigma-Aldrich) containing 0.65% agarose (Nacalai tesque, Kyoto, Japan), 2nM L-glutamine (Sanko Junyaku Co., Tokyo, Japan), 10mM HEPES (Dojindo, Kumamoto, Japan) and PS, and these were incubated at 37°C until the appropriate sized plaque were observed (approx. three days). Cells were fixed with 10% formaldehyde (Fujifilm Wako) and stained with crystal violet. For plaque purification of virus, without cell fixation, a well-isolated plaque was picked and resuspended in 100 μL DMEM(-) and stored at -70°C. Then, a 50 μL aliquot from the 100 μL plaque solution was expanded in HRT18 cells with DMEM(-) in a 10 cm dish, and then these viruses were harvested after the appropriate CPE was observed and was stored at -70°C as a master stock. 200 μL of master virus was expanded in the cells in an 18 cm dish and was used as a working wtBToV.

### RNA extraction, RT-PCR and sequencing

Viral RNA was extracted from 100–250 μL of the supernatant using TRIzol® LS Reagent (Thermo Fisher Scientific) according to the manufacturer’s instructions. Complementary DNA was synthesized using a SuperScript III® first-strand synthesis system (Thermo Fisher Scientific) and Oligo dT_15_ primer (TaKaRa, Shiga, Japan) or Oligo dT-3 sites Adaptor Primer for 3’ RACE (Takara). To analyze the complete BToV genome, seven fragments of about 3500–4500 base pair (bp) overlapping 170–360 bp (which correspond to BToV-derived sequence of fragments BToV-A to -G described below) and one fragment of about 320 bp of the 3’-terminal region by 3’ RACE were amplified using Phusion High-Fidelity PCR Kit (New England Biolabs, MA, USA) under the following conditions [98°C, 50 s; 35 cycles of (98°C, 10 s; 60–65°C, 30 s; 72°C, 1 min/kb); 72°C 10 min], and these PCR fragments were purified with AMPure XP (Beckman Coulter, CA, USA). 5’ ends of the viral genome were analyzed by 5’-Full RACE Core Set (TaKaRa) and the PCR products of 5’ RACE were cloned using pGEM®-T Easy Vector Systems (Promega, WI, USA). Purified PCR fragments and pGEM cloning 5’-terminal sequence of BToV were sequenced using primers designed for every ∼300 bp of the BToV genome. Sequences were assembled by MEGA6 (71) or Sequencher software (Hitachi High-Tech, Tokyo, Japan). Even without 5’ RACE, almost the entire full-length sequencing was achieved, except for 21 bp at the 5’ terminal end derived from the primer, which was applied to the full-length sequencing of rBToV and rEGFP c1.

### BAC construction of wtBToV

pBeloBAC11 containing CMV immediate-early promoter, Rz, and BGH termination (48) was used. Full-length cDNA of the BToV genome was assembled under the control of the CMV promoter and was flanked at the 3’-terminal end by a poly(A) tail (Fig. 1A), using a Red/ET recombination system counterselection BAC modification kit (Gene Bridges Heidelberg, Germany) according to the manufacturer’s instructions (Fig. 1B). Briefly, *E. coli* strain NEB10ß (Bio-Rad) carrying the BAC was electroporated with the pRed/ET expression plasmid using an Eppendorf Eporator® (Eppendorf, Hamburg, Germany). The Red/ET recombinant enzyme was induced by L-arabinose (Sigma-Aldrich) and a temperature shift from 30°C to 37°C, and then linear rpsL/Neo counter-selection/selection cassette (prepared by PCR) flanked by hms was electroporated so that the Red/ET recombination inserts the rpsL/Neo cassette into the target position through hms. Only *E. coli* carrying the BAC with the rpsL/Neo cassette could be selected by kanamycin at 30°C. The Red/ET enzyme was induced in the same way in *E. coli* carrying modified BAC and was electroporated with the linear cDNA fragment of BToV flanked by hms. Red/ET recombination could replace the rpsL/Neo cassette with the cDNA fragment of BToV. Only *E. coli* carrying the BAC with the target BToV cDNA fragment could be selected by streptomycin at 37°C at which the pRed/ET plasmid was removed (Fig. 1B). To use this system, the full-length genomic sequence of BToV was divided into eight fragments flanked by the hms and was assembled into the BAC plasmid in a sequential order (Fig. 1A). To distinguish the base numbers between BToV and BAC, BToV genome is 28,313 nt (without poly-A and was written as the normal number) while that of BAC is 8,219 bp, written as the number in parentheses starting from the CMV promoter. The BToV-derived sequences of each fragment correspond to nt 1 to 4,497 (BToV-A), nt 4,139 to 8,512 (BToV-B), nt 8,301 to 12,748 (BToV-C), nt 12,413 to 16,612 (BToV-D), nt 16,444 to 20,731(BToV-E), nt 20,512 to 24,877 (BToV-F), nt 24,675 to 28,196 (BToV-G), and nt 27,987 to 28,313 (BToV-H), respectively. The 5’ terminal hm of the BToV-A fragment and the other fragments were regions of a partial CMV promoter, (205) to (605), and regions overlapping between the fragments, respectively. The 3’ terminal hm of all fragments is the region including Rz and BGH, which is (606) to (1,001) (Fig. 1A). Fragments BToV-A to -G were prepared using standard PCR protocols with the overlap extension technique (72). Fragment BToV-H was prepared through PCR using a plasmid template containing a chemically synthesized BToV sequence between nt 27,987 to 28,313, 25 nt of adenine (pA), and the hm of the 3’ terminal sequence described above (Eurofins, Luxembourg, Luxembourg). All fragments were amplified using KOD Plus Neo (TOYOBO, Osaka, Japan) and purified with gel-extraction and phenol-chloroform extraction before electroporation. After assembly of the eight fragments, BAC carrying the cDNA clone of the full-length BToV (pBAC-BToV^mut1^) contained three mutations, namely, two single mutations at nt G3399T (E871D in NSP1 protein) and at nt T8469G (silent), and the *E. coli* chromosomal derived sequence of 1,350 bp was inserted at nt 10,136 (Fig. 2A). pRed/ET recombination method could revert two substitutions but failed to delete the *E. coli* chromosomal derived sequence. To delete this sequence in vitro, type IIS restriction enzyme SapI (New England Biolabs), which recognizes asymmetric DNA sequences and cleave outsides their 5’-GCTCTTCN↓NNN -3’ recognition site, was used. First, two SapI sites at nt (4,526) and (5,736) near or in the SopA gene of BAC plasmid were removed by introducing single mutations, 5’-(4,526) GCTCTTC → <u>C</u>CTCTTC and 5’-(5,736) GCTCTTC → G<u>G</u>TCTTC respectively, using NEBuilder HiFi DNA Assembly Master Mix (New England Biolabs) and two NgoMVI restriction sites located outside the two SapI sites at nt (3,633) and (7,549). Next, SapI recognition sites were inserted on both sides (one in the reverse direction) of the *E. coli* chromosomal derived sequence by pRed/ET recombination. Finally, the resulting BAC plasmid (pBAC-BToV^mut2^) was treated with SapI and was subjected to in vitro self-ligation (pBAC-BToV^pre^). pBAC-BToV^pre^ was electroporated into NEB10ß and was successfully propagated.

### BAC construction of mutant BToVs

To introduce mutations into the HE gene of pBAC-BToV^pre^, the rpsL/Neo cassette was inserted through the 5’ terminal hm including a partial M gene, nt 26,037 to 26,249 and the 3’ terminal hm including 3’-UTR, poly(A), Rz, and BGH, 28,150 to (1,001), using pRed/ET recombination (Fig. 2B). The resulting BAC with the rpsL/Neo cassette (pBAC-BToV^HE/N-rpsL/Neo^) allows manipulation of not only the HE gene but also the N gene. The rpsL/Neo cassette was replaced with the mutated HE gene or with the Enhanced Green Fluorescent Protein (EGFP) gene (Fig. 2B).

Cell-adapted wtBToV in our laboratory lost full-length HE (HEf) due to a stop codon at nt 481 (the base number of HE gene), CAG (Q)→<u>T</u>AG (stop), resulting in the soluble form HE 160 aa in length (HEs) (36, 37) (Fig. 3A). Using pBAC-BToV^HE/N-rpsL/Neo^, four pBAC plasmids were created; pBAC-BToV with two genetic markers at nt T207C (T26530C) and nt T228A (T26554A), pBAC-BToV-HEf with the replacement of a stop codon (TAG) to CAG to express full-length HE, pBAC-BToV-HEs/HA and pBAC-BToV-HEf/HA with the addition of HA-tag at the C-terminal end of HEs and HEf proteins, respectively. pBAC-BToV-HEf/HA had the HE gene mutation at nt 1,029, GAATTC→GAA<u>C</u>TC (silent) to remove the EcoRI site. In addition, pBAC-BToV-EGFP in which the HE gene was completely replaced with EGFP was created. All BAC plasmids were purified with NucleoBond Xtra Maxi (Macherey-Nagel, Dueren, Germany) from 1,000mL cultured *E. coli* cells.

### Rescue of recombinant BToVs

293T and HRT18 cells were seeded into 24-well plates at 1.5 × 10^5^ and 2.0 × 10^5^ cells/well, respectively, on the day before the experiment. 293T cells were transfected with 2.0 μg pBAC-BToV or -HE mutant and - EGFP with 1.5 μL Lipofectamine 3000 and 1.0 μL P3000 Regent (Thermo Fisher Scientific) and incubated at 37°C. The next day, DMEM(+) was replaced with 500 μL DMEM(-) to remove FBS. After 3 dpt, supernatants were harvested. The 50 μL supernatant was inoculated to susceptible HRT18 cells which had been washed with phosphate buffered saline (PBS) twice after which 450 μL DMEM (-) was added. After 2 dpi, supernatants were harvested and viruses were plaque-purified three times. A 20 μL aliquot from the 100 μL plaque suspension was added to HRT18 cells with 480 μL DMEM(-) in 24-well plates, and the supernatant was harvested after 2 dpi. This was stored at -70°C as a master virus. Next, 100 μL of the master viruses were expanded on HRT18 cells with 10 mL DMEM(-) in a 10 cm dish, and these viruses were harvested after the appropriate CPE was observed. These were stored as passage 0 (P0) viruses, which were used for subsequent experiments. In the case of rEGFP, plaque purification of some recombinant viruses (No.3 to No.6) were omitted because of EGFP gene instability. 293T cells prepared on a 3 cm dish were transfected with 5 times reagent volumes and pBAC-BToV-EGFP. The 2.5 mL supernatants of transfected HRT18 cells were inoculated into the cells in a 10 cm dish to expand the viruses, and these viruses were then harvested. In this case, these were used as passage 0 (P0) viruses. The nomenclature of recombinant BToV rescued from these BAC plasmids was rBToV(rHEs), rHEf, rHEs/HA, rHEf/HA and rEGFP, respectively (Fig. 3A).

### Immunofluorescence staining

A 35 mm glass base dish (12 mm glass area) (Iwaki, Shizuoka, Japan) was seeded with 1.0 × 10^5^ HRT18 cells, the day before the experiment. Cells were washed twice with PBS and infected with 300 μL recombinant BToV in DMEM(-) with a MOI of 0.05. After 24 hpi (and 36 hpi only for the rEGFP), infected cells were washed with PBS, fixed with 4% paraformaldehyde for 20 min at room temperature (rt), and were permeabilized with 0.2% Triton X-100 for 20 min at RT. Cells were incubated with mouse anti-HE, anti-M antiserum, or rabbit and mouse anti-HA antibodies at 4°C overnight, and were then incubated with a mixed FITC-conjugated goat anti-mouse or mixed AlexaFluor™ 594-conjugated goat anti-mouse for mouse antiserum and FITC-conjugated goat anti-rabbit antibodies for rabbit antibody, for 2 h at rt. Cell nuclei were stained with Hoechst 33258 solution (Dojindo) for 20 min at rt. PBS washing was done twice between each step. Stained cells were observed under confocal laser scanning microscopy using an LSM 710 laser scanning microscope (Carl Zeiss, Oberkochen, Germany).

### α-naphthyl acetate esterase activity

HRT18 cells grown in 24-well plates were infected with recombinant BToVs with a MOI of 0.05. At 24 hpi, cells were fixed with fixative solution, stained for a α-naphthyl acetate esterase with a α-NA esterase staining kit (Muto Pure Chemicals, Tokyo, Japan) according to the manufacturer’s instructions, and observed under phase-contrast microscopy.

### Immunoblotting

HRT18 cells grown in 24-well plates were infected with the recombinant BToVs with a MOI of 0.05. COS7 cells were transfected with 1.0 μg pCAGGS-HEf or pCAGGS-HEs with 1.5 μL Lipofectamine 3000 and 2.0 μl P3000 Regent. At 24 hpi (and 36 hpi only for rEGFP), infected HRT18 cells and 40 hpt transfected COS7 cells were lysed with 100 μL sample buffer (50 mM Tris, 2% SDS, 0.1% bromophenol blue, 10% glycerol, and 1% 2-mercaptoethanol) and were boiled for 5 min. N protein, EGFP, and HA-tagged proteins of samples were analyzed by sodium dodecyl sulfate-polyacrylamide gel electrophoresis (SDS-PAGE) and were transferred to a PVDF membrane (Merck Millipore, Tokyo, Japan). The membrane was incubated with rabbit anti-N antiserum or mouse anti-GFP antibody, and then with HRP-conjugated anti-rabbit and anti-mouse antibodies, respectively. To detect HA-tagged proteins, mouse-anti-HA 12CA5 antibody for HA-tagged HEf and rabbit anti-HA antibody for HA-tagged HEs were used because of differences in their reactivity. Protein bands were visualized with ECL Prime Western Blotting Detection Reagent (GE Healthcare, IL, USA) on a Light Capture II instrument (ATTO, Tokyo, Japan).

### Growth kinetics of recombinant BToVs

HRT18 cells in 24-well plates were washed twice with PBS and were infected with recombinant BToVs with a MOI of 0.001. After 1.5 h adsorption at 37°C, unadsorbed viruses were removed and cells were washed twice with PBS, and 0.5 mL DMEM(-) was added. Viruses in the supernatant were collected at 2, 24, 48, and 72 hpi and the supernatant was clarified by centrifugation at 12,000 rpm for 5 min at 4°C. Virus titers in the culture media were determined in a 96-well plate by TCID_50_, as previously described (73).

### Measurement of the stability of HE and EGFP gene

HRT18 cells in a 24-well plate were washed twice with PBS after which 0.5 mL DMEM(-) was added; these were then infected with recombinant BToVs with a MOI of 0.001. After the appropriate CPE was observed (2 to 3 dpi), the supernatant was harvested and 0.5 uL was inoculated to the fresh cells, resulting in a serial passages by 1,000-fold diluted virus. To estimate stability of the HEf and EGFP genes during serial passages, HRT18 cells in a 24-well plate were infected with rHEf and rHEf/HA harvested at the indicated passage history with a MOI of 0.05, and cells at 24 hpi were subjected to an α-NA esterase assay or to immunoblotting of N proteins. EGFP-expression of the infected cells with rEGFP were monitored during serial passages under a fluorescence microscope (Olympus CKX41 U-RFLT50; Olympus, Tokyo, Japan). Later, images of cells infected with rEGFP harvested at the indicated passage history at the same conditions were taken, and EGFP and N proteins were detected by immunoblotting. Supernatants of the infected cells (resulting in +1 passage history) were subjected to RNA extraction, RT-PCR, and sequencing described above. PCR fragments of 1130 and 1630 bp containing HE and EGFP genes, respectively, were amplified using Takara ExTaq (Takara) under the following conditions [94°C, 5min; 30 cycles of (94°C, 30 s; 50°C, 30 s; 72°C, 1 min/kb); 72°C 10min]. DNase treatment was also performed before RNA extraction to digest BAC plasmid from transfection, depending on passage history.

## Acknowledgments

We thank Dr. Tsunemitsu Hiroshi (Nishimikawa Livestock Hygiene Service Center, Japan) for providing cell-adapted BToV (Aichi strain) and Drs. Asanuma Hideki (National Institute of Infectious Diseases, Japan) and Yuasa Noriyuki (Tokyo chemical industry co. Ltd., Japan) for providing mouse anti-HE and anti-M antiserum, respectively. This work was supported by a Grant-in-Aid for Scientific Research (C; No.19K06393) from the Ministry of Education, Culture, Sports, Science and Technology in Japan, and by the Grant for Joint Research Project of the Research Institute for Microbial Diseases, Osaka University.

## References

1. ICTV (International Committee on Taxonomy of Virus). 2019. Virus Taxonomy: 2019 Release.

2. Vanopdenbosch E, Wellemans G, Petroff K. 1991. Breda virus associated with respiratory disease in calves. Vet Rec 129:203.

3. Duckmanton L, Luan B, Devenish J, Tellier R, Petric M. 1997. Characterization of torovirus from human fecal specimens. Virology 239:158–68.

4. Kroneman A, Cornelissen LA, Horzinek MC, de Groot RJ, Egberink HF. 1998. Identification and characterization of a porcine torovirus. J Virol 72:3507–11.

5. Woode GN, Reed DE, Runnels PL, Herrig MA, Hill HT. 1982. Studies with an unclassified virus isolated from diarrheic calves. Vet Microbiol 7:221–40.

6. Weiss M, Steck F, Horzinek MC. 1983. Purification and Partial Characterization of a New Enveloped RNA Virus (Berne Virus). J Gen Virol 64:1849–1858.

7. Ito T, Okada N, Okawa M, Fukuyama S, Shimizu M. 2009. Detection and characterization of bovine torovirus from the respiratory tract in Japanese cattle. Vet Microbiol 136:366–371.

8. SH L, HY K, EW C, D K. 2019. Causative Agents and Epidemiology of Diarrhea in Korean Native Calves. J Vet Sci 20.

9. Nogueira JS, Asano KM, de Souza SP, Brand ão PE, Richtzenhain LJ. 2013. First detection and molecular diversity of Brazilian bovine torovirus (BToV) strains from young and adult cattle. Res Vet Sci 95:799– 801.

10. Lojkić I, Krešić N, Šimić I, Bedeković T. 2015. Detection and molecular characterisation of bovine corona and toroviruses from Croatian cattle. BMC Vet Res 11:202.

11. Koopmans M, van Wuijckhuise-Sjouke L, Schukken YH, Cremers H, Horzinek MC. 1991. Association of diarrhea in cattle with torovirus infections on farms. Am J Vet Res 52:1769–73.

12. Hoet AE, Saif LJ. 2004. Bovine torovirus (Breda virus) revisited. Anim Heal Res Rev 5:157–71.

13. Hoet AE, Nielsen PR, Hasoksuz M, Thomas C, Wittum TE, Saif LJ. 2003. Detection of Bovine Torovirus and other Enteric Pathogens in Feces from Diarrhea Cases in Cattle. J Vet Diagnostic Investig 15:205– 212.

14. Gülaçtiİ, I ą F; idan H, Sözdutmazİ. 2014. Detection of bovine torovirus in fecal specimens from calves with diarrhea in Turkey. Arch Virol 159:1623–1627.

15. Duckmanton L, Carman S, Nagy E, Petric M. 1998. Detection of bovine torovirus in fecal specimens of calves with diarrhea from Ontario farms. J Clin Microbiol 36:1266–70.

16. H L, B Z, H Y, C T. 2020. First Detection and Genomic Characteristics of Bovine Torovirus in Dairy Calves in China. Arch Virol 165.

17. Zhou L, Wei H, Zhou Y, Xu Z, Zhu L, Horne J. 2014. Molecular epidemiology of Porcine torovirus (PToV) in Sichuan Province, China: 2011–2013. Virol J 11:106.

18. Y F, Y K, F S, H A, R I, K S, Y K, T O, M O, T F, S T, Y O, J S, T M, T O, M N. 2020. Complete Genome Sequencing and Genetic Analysis of a Japanese Porcine Torovirus Strain Detected in Swine Feces. Arch Virol 165.

19. Shin D-J, Park S-I, Jeong Y-J, Hosmillo M, Kim H-H, Kim H-J, Kwon H-J, Kang M-I, Park S-J, Cho K-O. 2010. Detection and molecular characterization of porcine toroviruses in Korea. Arch Virol 155:417–422.

20. Pignatelli J, Grau-Roma L, Jiménez M, Segalés J, Rodríguez D. 2010. Longitudinal serological and virological study on porcine torovirus (PToV) in piglets from Spanish farms. Vet Microbiol 146:260–268.

21. Anbalagan S, Peterson J, Wassman B, Elston J, Schwartz K. 2014. Genome sequence of torovirus identified from a pig with porcine epidemic diarrhea virus from the United States. Genome Announc 2.

22. ZM H, YL Y, LD X, B W, P Q, YW H. 2019. Porcine Torovirus (PToV)-A Brief Review of Etiology, Diagnostic Assays and Current Epidemiology. Front Vet Sci 6.

23. SL S, EJ S, RJ de G. 2006. Characterization of a Torovirus Main Proteinase. J Virol 80.

24. Snijder EJ, Horzinek MC. 1993. Toroviruses: replication, evolution and comparison with other members of the coronavirus-like superfamily. J Gen Virol 74:2305–2316.

25. Draker R, Roper RL, Petric M, Tellier R. 2006. The complete sequence of the bovine torovirus genome. Virus Res 115:56–68.

26. Snijder EJ, Den Boon JA, Spaan WJM, Verjans GMGM, Horzinek MC. 1989. Identification and Primary Structure of the Gene Encoding the Berne Virus Nucleocapsid Protein. J Gen Virol 70:3363–3370.

27. Horzinek MC, Ederveen J, Weiss M. 1985. The Nucleocapsid of Berne Virus. J Gen Virol 66:1287–1296.

28. Den Boon JA, Snijder EJ, Locker JK, Horzinek MC, Rottier PJ. 1991. Another triple-spanning envelope protein among intracellularly budding RNA viruses: the torovirus E protein. Virology 182:655–63.

29. Snijder EJ, Den Boon JA, Spaan WJ, Weiss M, Horzinek MC. 1990. Primary structure and post-translational processing of the Berne virus peplomer protein. Virology 178:355–63.

30. Cornelissen LA, Wierda CM, van der Meer FJ, Herrewegh AA, Horzinek MC, Egberink HF, de Groot RJ. 1997. Hemagglutinin-esterase, a novel structural protein of torovirus. J Virol 71:5277–86.

31. Smits SL, Lavazza A, Matiz K, Horzinek MC, Koopmans MP, de Groot RJ. 2003. Phylogenetic and evolutionary relationships among torovirus field variants: evidence for multiple intertypic recombination events. J Virol 77:9567–77.

32. Cong Y, Zarlenga DS, Richt JA, Wang X, Wang Y, Suo S, Wang J, Ren Y, Ren X. 2013. Evolution and homologous recombination of the hemagglutinin–esterase gene sequences from porcine torovirus. Virus Genes 47:66–74.

33. Ito M, Tsuchiaka S, Naoi Y, Otomaru K, Sato M, Masuda T, Haga K, Oka T, Yamasato H, Omatsu T, Sugimura S, Aoki H, Furuya T, Katayama Y, Oba M, Shirai J, Katayama K, Mizutani T, Nagai M. 2016. Whole genome analysis of Japanese bovine toroviruses reveals natural recombination between porcine and bovine toroviruses. Infect Genet Evol 38:90–95.

34. Aita T, Kuwabara M, Murayama K, Sasagawa Y, Yabe S, Higuchi R, Tamura T, Miyazaki A, Tsunemitsu H. 2012. Characterization of epidemic diarrhea outbreaks associated with bovine torovirus in adult cows. Arch Virol 157:423–431.

35. Ito T, Okada N, Fukuyama S. 2007. Epidemiological analysis of bovine torovirus in Japan. Virus Res 126:32–37.

36. Kuwabara M, Wada K, Maeda Y, Miyazaki A, Tsunemitsu H. 2007. First Isolation of Cytopathogenic Bovine Torovirus in Cell Culture from a Calf with Diarrhea. Clin Vaccine Immunol 14:998–1004.

37. Shimabukuro K, Ujike M, Ito T, Tsunemitsu H, Oshitani H, Taguchi F. 2013. Hemagglutination mediated by the spike protein of cell-adapted bovine torovirus. Arch Virol 158:1561.

38. Weiss M, Horzinek MC. 1986. Morphogenesis of Berne Virus (Proposed Family Toroviridae). J Gen Virol 67:1305–1314.

39. Horzinek MC, Weiss M, Ederveen J. 1984. Berne Virus is Not “Coronavirus-like.” J Gen Virol 65:645–649.

40. Fagerland JA, Pohlenz JFL, Woode GN. 1986. A Morphological Study of the Replication of Breda Virus (Proposed Family Toroviridae) in Bovine Intestinal Cells. J Gen Virol 67:1293–1304.

41. Corman VM, Muth D, Niemeyer D, Drosten C. 2018. Hosts and Sources of Endemic Human Coronaviruses. Adv Virus Res 100:163–188.

42. Drosten C, Gunther S, Preiser W, van der Werf S, Brodt HR, Becker S, Rabenau H, Panning M, Kolesnikova L, Fouchier RA, Berger A, Burguiere AM, Cinatl J, Eickmann M, Escriou N, Grywna K, Kramme S, Manuguerra JC, Muller S, Rickerts V, Sturmer M, Vieth S, Klenk HD, Osterhaus AD, Schmitz H, Doerr HW. 2003. Identification of a novel coronavirus in patients with severe acute respiratory syndrome. N Engl J Med2003/04/12. 348:1967–1976.

43. Kuiken T, Fouchier RA, Schutten M, Rimmelzwaan GF, van Amerongen G, van Riel D, Laman JD, de Jong T, van Doornum G, Lim W, Ling AE, Chan PK, Tam JS, Zambon MC, Gopal R, Drosten C, van der Werf S, Escriou N, Manuguerra JC, Stohr K, Peiris JS, Osterhaus AD. 2003. Newly discovered coronavirus as the primary cause of severe acute respiratory syndrome. Lancet2003/08/02. 362:263–270.

44. Zaki AM, van Boheemen S, Bestebroer TM, Osterhaus AD, Fouchier RA. 2012. Isolation of a novel coronavirus from a man with pneumonia in Saudi Arabia. N Engl J Med2012/10/19. 367:1814–1820.

45. Zhou P, Yang X-L, Wang X-G, Hu B, Zhang L, Zhang W, Si H-R, Zhu Y, Li B, Huang C-L, Chen H-D, Chen J, Luo Y, Guo H, Jiang R-D, Liu M-Q, Chen Y, Shen X-R, Wang X, Zheng X-S, Zhao K, Chen Q-J, Deng F, Liu L-L, Yan B, Zhan F-X, Wang Y-Y, Xiao G-F, Shi Z-L. 2020. A pneumonia outbreak associated with a new coronavirus of probable bat origin. Nature 579:270–273.

46. R L, X Z, J L, P N, B Y, H W, W W, H S, B H, N Z, Y B, X M, F Z, L W, T H, H Z, Z H, W Z, L Z, J C, Y M, J W, Y L, J Y, Z X, J M, WJ L, D W, W X, EC H, GF G, G W, W C, W S, W T. 2020. Genomic Characterisation and Epidemiology of 2019 Novel Coronavirus: Implications for Virus Origins and Receptor Binding. Lancet (London, England) 395.

47. Yount B, Curtis KM, Baric RS. 2000. Strategy for systematic assembly of large RNA and DNA genomes: transmissible gastroenteritis virus model. J Virol 74:10600–11.

48. F A, JM G, Z P, A I, E C, J P-D, L E. 2000. Engineering the Largest RNA Virus Genome as an Infectious Bacterial Artificial Chromosome. Proc Natl Acad Sci U S A 97:5516–21.

49. G T, R H-L, I S, V T, HJ T. 2008. Genome Organization and Reverse Genetic Analysis of a Type I Feline Coronavirus. J Virol 82.

50. Donaldson EF, Yount B, Sims AC, Burkett S, Pickles RJ, Baric RS. 2008. Systematic assembly of a full-length infectious clone of human coronavirus NL63. J Virol 82:11948–57.

51. Á B, A F, Z Z, Á H, L D, F A, L E, S B. 2012. Molecular Characterization of Feline Infectious Peritonitis Virus Strain DF-2 and Studies of the Role of ORF3abc in Viral Cell Tropism. J Virol 86.

52. Tekes G, Spies D, Bank-Wolf B, Thiel V, Thiel H-J. 2012. A reverse genetics approach to study feline infectious peritonitis. J Virol 86:6994–8.

53. F A, ML D, I S, S Z, JL N-T, S M-J, G A, L E. 2013. Engineering a Replication-Competent, Propagation-Defective Middle East Respiratory Syndrome Coronavirus as a Vaccine Candidate. MBio 4:e00650–13.

54. Scobey T, Yount BL, Sims AC, Donaldson EF, Agnihothram SS, Menachery VD, Graham RL, Swanstrom J, Bove PF, Kim JD, Grego S, Randell SH, Baric RS. 2013. Reverse genetics with a full-length infectious cDNA of the Middle East respiratory syndrome coronavirus. Proc Natl Acad Sci U S A 110:16157–62.

55. Beall A, Yount B, Lin C-M, Hou Y, Wang Q, Saif L, Baric R. 2016. Characterization of a Pathogenic Full-Length cDNA Clone and Transmission Model for Porcine Epidemic Diarrhea Virus Strain PC22A. MBio 7:e01451–15.

56. Ehmann R, Kristen-Burmann C, Bank-Wolf B, König M, Herden C, Hain T, Thiel H-J, Ziebuhr J, Tekes G. 2018. Reverse Genetics for Type I Feline Coronavirus Field Isolate To Study the Molecular Pathogenesis of Feline Infectious Peritonitis. MBio 9.

57. D M, B M, D N, S S, N O, MA M, C D. 2017. Transgene Expression in the Genome of Middle East Respiratory Syndrome Coronavirus Based on a Novel Reverse Genetics System Utilizing Red-mediated Recombination Cloning. J Gen Virol 98.

58. Terada Y, Kuroda Y, Morikawa S, Matsuura Y, Maeda K, Kamitani W. 2019. Establishment of a Virulent Full-Length cDNA Clone for Type I Feline Coronavirus Strain C3663. J Virol 93.

59. Thiel V, Herold J, Schelle B, Siddell SG. 2001. Infectious RNA transcribed in vitro from a cDNA copy of the human coronavirus genome cloned in vaccinia virus. J Gen Virol 82:1273–1281.

60. T TNT, F L, N E, P V, H S, J P, J K, S S, M H, A K, M G, K S, L L, L H, M W, S P, D H, V C, S C-P, S S, D M, D N, VM C, MA M, C D, R D, J J, V T. 2020. Rapid Reconstruction of SARS-CoV-2 Using a Synthetic Genomics Platform. Nature 582:561–565.

61. Hou YJ, Okuda K, Edwards CE, Martinez DR, Asakura T, Dinnon KH, Kato T, Lee RE, Yount BL, Mascenik TM, Chen G, Olivier KN, Ghio A, Tse L V., Leist SR, Gralinski LE,Schäfer A, Dang H, Gilmore R, Nakano S, Sun L, Fulcher ML, Livraghi-Butrico A, Nicely NI, Cameron M, Cameron C, Kelvin DJ, de Silva A, Margolis DM, Markmann A, Bartelt L, Zumwalt R, Martinez FJ, Salvatore SP, Borczuk A, Tata PR, Sontake V, Kimple A, Jaspers I, O’Neal WK, Randell SH, Boucher RC, Baric RS. 2020. SARS-CoV-2 Reverse Genetics Reveals a Variable Infection Gradient in the Respiratory Tract. Cell.

62. Casais R, Thiel V, Siddell SG, Cavanagh D, Britton P. 2001. Reverse Genetics System for the Avian Coronavirus Infectious Bronchitis Virus. J Virol 75:12359–12369.

63. B Y, MR D, SR W, RS B. 2002. Systematic Assembly of a Full-Length Infectious cDNA of Mouse Hepatitis Virus Strain A59. J Virol 76:11065–78.

64. B Y, KM C, EA F, LE H, PB J, E P, MR D, TW G, RS B. 2003. Reverse Genetics With a Full-Length Infectious cDNA of Severe Acute Respiratory Syndrome Coronavirus. Proc Natl Acad Sci U S A 100:12995–3000.

65. Coley SE, Lavi E, Sawicki SG, Fu L, Schelle B, Karl N, Siddell SG, Thiel V. 2005. Recombinant mouse hepatitis virus strain A59 from cloned, full-length cDNA replicates to high titers in vitro and is fully pathogenic in vivo. J Virol 79:3097–106.

66. F A, ML D, C G, D E, E A, J O, I S, S Z, S A, JL M, A N, C C, L E. 2006. Construction of a Severe Acute Respiratory Syndrome Coronavirus Infectious cDNA Clone and a Replicon to Study Coronavirus RNA Synthesis. J Virol 80:10900–6.

67. St-Jean JR, Desforges M, Almazán F, Jacomy H, Enjuanes L, Talbot PJ. 2006. Recovery of a neurovirulent human coronavirus OC43 from an infectious cDNA clone. J Virol 80:3670–4.

68. MM B, RL G, EF D, B R, AC S, T S, RJ P, D C, RE J, RS B, MR D. 2008. Synthetic Recombinant Bat SARS-like Coronavirus Is Infectious in Cultured Cells and in Mice. Proc Natl Acad Sci U S A 105.

69. Ito T, Katayama S, Okada N, Masubuchi K, Fukuyama S i., Shimizu M. 2010. Genetic and Antigenic Characterization of Newly Isolated Bovine Toroviruses from Japanese Cattle. J Clin Microbiol 48:1795– 1800.

70. Kadowaki S, Chen Z, Asanuma H, Aizawa C, Kurata T, Tamura S. 2000. Protection against influenza virus infection in mice immunized by administration of hemagglutinin-expressing DNAs with electroporation. Vaccine 18:2779–88.

71. K T, G S, D P, A F, S K. 2013. MEGA6: Molecular Evolutionary Genetics Analysis Version 6.0. Mol Biol Evol 30.

72. Horton MR, Pease LR. 1991. Recombination and mutagenesis of DNA-sequences using PCR. In Directed mutagenesis: a practical approach. Edited by M. J. McPherson. 217–247.

73. Reed LJ, Muench H. 1938. A SIMPLE METHOD OF ESTIMATING FIFTY PER CENT ENDPOINTS12. Am J Epidemiol 27:493–497.

74. van Vliet Alw, Smits SL, Rottier PJM, Groot RJ de. 2002. Discontinuous and non-discontinuous subgenomic RNA transcription in a nidovirus. EMBO J 21:6571–6580.

75. Suzuki T, Terada Y, Enjuanes L, Ohashi S, Kamitani W. 2018. S1 Subunit of Spike Protein from a Current Highly Virulent Porcine Epidemic Diarrhea Virus Is an Important Determinant of Virulence in Piglets. Viruses 10.

76. Terada Y, Kawachi K, Matsuura Y, Kamitani W. 2017. MERS coronavirus nsp1 participates in an efficient propagation through a specific interaction with viral RNA. Virology 511:95–105.

77. Sakai Y, Kawachi K, Terada Y, Omori H, Matsuura Y, Kamitani W. 2017. Two-amino acids change in the nsp4 of SARS coronavirus abolishes viral replication. Virology 510:165–174.

78. Muth D, Meyer B, Niemeyer D, Schroeder S, Osterrieder N, Müller MA, Drosten C. 2017. Transgene expression in the genome of Middle East respiratory syndrome coronavirus based on a novel reverse genetics system utilizing Red-mediated recombination cloning. J Gen Virol 98:2461–2469.

79. De Groot RJ. 2006. Structure, function and evolution of the hemagglutinin-esterase proteins of corona-and toroviruses. Glycoconj J 23:59–72.

80. B S, HJ G, R B, HD K, G H. 1990. Hemagglutinating encephalomyelitis virus attaches to N-acetyl-9-O-acetylneuraminic acid-containing receptors on erythrocytes: comparison with bovine coronavirus and influenza C virus. Virus Res 16.

81. Vlasak R, Luytjes W, Spaan W, Palese P. 1988. Human and bovine coronaviruses recognize sialic acid-containing receptors similar to those of influenza C viruses. Proc Natl Acad Sci 85:4526–4529.

82. Desforges M, Desjardins J, Zhang C, Talbot PJ. 2013. The Acetyl-Esterase Activity of the Hemagglutinin-Esterase Protein of Human Coronavirus OC43 Strongly Enhances the Production of Infectious Virus. J Virol 87:3097–3107.

83. Bakkers MJG, Lang Y, Feitsma LJ, Hulswit RJG, de Poot Sah, van Vliet Alw, Margine I, de Groot-Mijnes Jdf, van Kuppeveld Fjm, Langereis MA, Huizinga EG, de Groot RJ. 2017. Betacoronavirus Adaptation to Humans Involved Progressive Loss of Hemagglutinin-Esterase Lectin Activity. Cell Host Microbe 21:356–366.

84. Williams RK, Jiang GS, Holmes K V. 1991. Receptor for mouse hepatitis virus is a member of the carcinoembryonic antigen family of glycoproteins. Proc Natl Acad Sci 88:5533–5536.

85. Langereis MA, Vliet ALW van, Boot W, Groot RJ de. 2010. Attachment of Mouse Hepatitis Virus to O-Acetylated Sialic Acid Is Mediated by Hemagglutinin-Esterase and Not by the Spike Protein. J Virol 84:8970–8974.

86. Luytjes W, Bredenbeek PJ, Noten AFH, Horzinek MC, Spaan WJM. 1988. Sequence of mouse hepatitis virus A59 mRNA 2: Indications for RNA recombination between coronaviruses and influenza C virus. Virology 166:415–422.

87. Yokomori K, Banner LR, Lai MMC. 1991. Heterogeneity of gene expression of the hemagglutinin-esterase (HE) protein of murine coronaviruses. Virology 183:647.

88. Lissenberg A, Vrolijk MM, Vliet ALW van, Langereis MA, Groot-Mijnes JDF de, Rottier PJM, Groot RJ de. 2005. Luxury at a Cost? Recombinant Mouse Hepatitis Viruses Expressing the Accessory Hemagglutinin Esterase Protein Display Reduced Fitness In Vitro. J Virol 79:15054–15063.

89. Lehmann KC, Snijder EJ, Posthuma CC, Gorbalenya AE. 2015. What we know but do not understand about nidovirus helicases. Virus Res 202:12–32.

90. Hao W, Wojdyla JA, Zhao R, Han R, Das R, Zlatev I, Manoharan M, Wang M, Cui S. 2017. Crystal structure of Middle East respiratory syndrome coronavirus helicase. PLoS Pathog 13:e1006474.

91. Tamura K, Stecher G, Peterson D, Filipski A, Kumar S. 2013. MEGA6: Molecular Evolutionary Genetics Analysis version 6.0. Mol Biol Evol 30:2725–9.

92. van Dinten LC, van Tol H, Gorbalenya AE, Snijder EJ. 2000. The predicted metal-binding region of the arterivirus helicase protein is involved in subgenomic mRNA synthesis, genome replication, and virion biogenesis. J Virol 74:5213–23.

93. Seybert A, Posthuma CC, van Dinten LC, Snijder EJ, Gorbalenya AE, Ziebuhr J. 2005. A complex zinc finger controls the enzymatic activities of nidovirus helicases. J Virol 79:696–704.

94. van Vliet Alw, Smits SL, Rottier PJM, de Groot RJ. 2002. Discontinuous and non-discontinuous subgenomic RNA transcription in a nidovirus. EMBO J 21:6571–80.

95. Stewart H, Brown K, Dinan AM, Irigoyen N, Snijder EJ, Firth AE. 2018. Transcriptional and Translational Landscape of Equine Torovirus. J Virol 92.

